# Evaluating goodness-of-fit of animal movement models using lineups

**DOI:** 10.1101/2023.09.26.559591

**Authors:** John Fieberg, Smith Freeman, Johannes Signer

## Abstract

Models of animal movement are frequently fit to animal location data to understand how animals respond to and interact with local environmental features. Several open-source software packages are available for analyzing animal movements and can facilitate parameter estimation, yet there are relatively few methods available for evaluating model goodness-of-fit. We describe how a simple graphical technique, the *lineup protocol*, can be used to evaluate goodness-of-fit of integrated step-selection analyses and hidden Markov models, but the method can be applied much more broadly. We leverage the ability to simulate data from fitted models, and demonstrate the approach using both methods applied to fisher (*Pekania pennanti*) data. A variety of responses and movement metrics can be used to evaluate models, and the lineup protocol can be tailored to focus on specific model assumptions or movement features that are of primary interest. Although it is possible to evaluate goodness-of-fit using a formal hypothesis test, the method can also be used in a more exploratory fashion (e.g., to visualize variability in model behavior across stochastic simulations or identify areas where the model could be improved). We provide coded examples and two vignettes to demonstrate the flexibility of the approach and encourage movement ecologists to consider how their models will be applied when choosing appropriate graphical responses for evaluating goodness-of-fit.

## 1 Introduction

Technological advances, including smaller and better tracking devices (Kays et al. 2015), have led to an exponential increase in animal location data, and have fueled the development of new statistical methods and software for modeling animal movement (Hooten et al. 2017, Joo et al. 2020). Integrated step-selection analyses (ISSAs) (Avgar et al. 2016, Fieberg et al. 2021) and hidden Markov models (HMMs) (Langrock et al. 2012, McClintock et al. 2012), in particular, are extremely popular for analyzing wildlife telemetry data due to the availability of open-source software (`amt and momentuHMM`) for implementing them (McClintock and Michelot 2018, Signer et al. 2019). These approaches, as well as recently developed methods that make it possible to combine the two frameworks (Klappstein et al. 2023, Pohle et al. 2023), allow researchers to fit rich models to telemetry data in which movements may vary spatially and temporally as a function of environmental features (e.g., landcover types, distance to roads) and latent (unobserved) behavioral states (e.g., that may be associated with whether the individual has recently fed).

Despite the popularity of these analytical frameworks, there are relatively few methods available for evaluating their goodness-of-fit. The `momentuHMM` package returns pseudo-residuals from fitted HMMs, which can be used to evaluate fit (e.g., using a quantile-quantile plot) or to explore potential violations of important assumptions (e.g., lack of autocorrelation in the residuals). Yet, many users are likely to struggle to interpret pseudo-residuals. Methods for validating ISSAs are almost non-existent, likely due to the “tricks” commonly used for inference; for example, a two-step procedure is typically used to estimate movement parameters via importance sampling (Avgar et al. 2016, Fieberg et al. 2021, Michelot et al. 2023), and random-effect specifications typically rely on a likelihood equivalence between Poisson regression with stratum-specific intercepts and conditional logistic regression models (Muff et al. 2020). One approach to model evaluation for ISSAs, recently implemented in the amt package and referred to as *used-habitat-calibration plots* (Fieberg et al. 2018), focuses on whether the distribution of environmental characteristics (e.g., elevation, landcover class) at model-predicted locations matches the distribution of those same characteristics measured at locations visited by the individual(s). This method relies on predictive simulation techniques to derive a simulation envelope for the distribution of environmental conditions at used locations, and shares similarities with Bayesian approaches that evaluate goodness-of-fit using posterior-predictive distributions (Conn et al. 2018).

Our goal here is to develop a more general and flexible approach for evaluating goodness-of-fit of ISSAs and HMMs, and animal movement models in general, by leveraging our ability to simulate trajectories from fitted models. Whereas Fieberg et al. (2018) derived predictive distributions using cross-validation at the step level (i.e., predicted locations were conditioned on the previously observed location), we will consider simulations of multi-step trajectories. We can then consider how these trajectories compare to observed movements using a wide range of statistical and graphical summaries. A rigorous goodness-of-fit test can be implemented using the lineup protocol, a formal graphical null hypothesis testing framework (Buja et al. 2009, Wickham et al. 2010). Alternatively, we can use the visualizations in an exploratory way to better understand the diversity of movements that may be generated from fitted models or to determine if our models are capable of generating important behaviors seen in real data.

## 2 The Lineup Protocol

As with any goodness-of-fit test, we will assume the following null and alternative hypotheses:

*H*_0_ : The data are consistent with the assumed model

*H_A_* : The data are not consistent with the assumed model

Although the lineup protocol takes its name from police lineups, the method actually depends on the power of the human eye to decipher patterns rather than to identify a known entity among unknown entities (Wickham et al. 2010). In short, we will simulate several data sets from our fitted model, summarize the simulated and observed data sets graphically, and see if the observed data are easily decipherable from simulated data sets. If they are, then we will reject the null hypothesis and conclude that our model is missing one or more important features. Otherwise, we will fail to reject the null hypothesis and conclude that the model successfully captures the characteristics of the observed data.

When using a lineup, a plot serves as our test statistic, and plots of the simulated data collectively serve as our sampling distribution under the null hypothesis. In our examples, we will generate multi-panel plots summarizing the observed data and 19 simulated data sets, with the observed-data panel randomly located (e.g., Figure 1). If the real data are consistent with the assumed model (i.e., if the null hypothesis is true), then we would have a 1 in 20 (or 0.05) chance of selecting any one of the panels. Thus, selecting the panel with the observed data implies that the p-value for the hypothesis test is *≤* 0.05; the *≤* reflects the fact that the precision with which we are able to calculate the p-value is limited by the number of panels included in the plot (we can increase precision by including more panels). To increase the statistical power of the test, one can offer lineups to multiple observers. In the simplest case, where a unique lineup is shown to each observer, the p-value for the goodness-of-fit test can be determined using the cumulative distribution function of the binomial distribution with *N* = the number of lineups/observers and *p* = 1*/n* (where *n* is the number of panels in each lineup). For example, if a unique lineup with 20 plots were shown to *N* = 10 independent observers, and 3 identified the panel containing the real data, the p-value would be given by 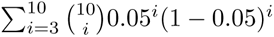 and could be calculated in R using `1 - pbinom(3, size = 10, p = 1/20`)) = 0.001.

**Figure 1:**
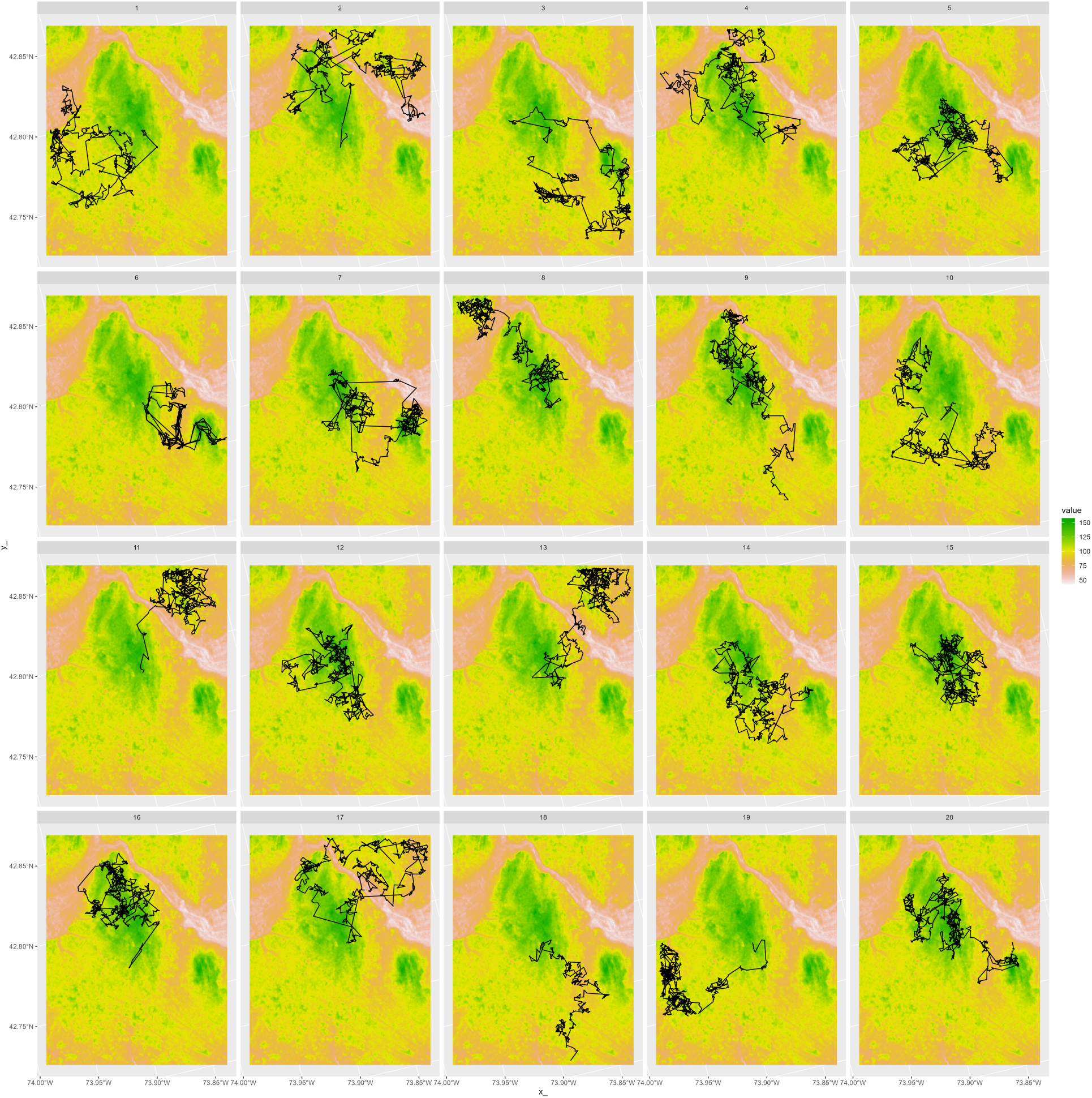
Simulated trajectories overlayed on a map of elevation. Nineteen of the panels were generated by simulating observations from a model of animal movement. The model was parameterized using an integrated step-selection analysis with fisher data from LaPoint et al. (2013). The other panel contains an observed trajectory for an individual from the same study but whose data were not used to parameterize the model.

## 3 Integrated Step-Selection Analysis of Fisher Data

To illustrate the lineup protocol, we first consider an ISSA fit to tracking data from 3 fishers (*Pekania pennanti*) available through Movebank (Brown et al. 2012, LaPoint et al. 2013a, LaPoint et al. 2013b). We resampled the data to a fix interval of 10 minutes with a tolerance of 1 minute using the `track_resample` function in the `R-package amt` (Signer et al. 2019). We pooled data from 2 of the individuals and set aside data from the 3rd individual for model evaluation.

We assumed that the distributions of step lengths and turn angles followed Gamma and von Mises distributions, respectively. We estimated tentative parameters of these distributions using the empirical steps connecting consecutive locations, from which we then simulated 5 random steps for each observed step using the `random_steps` function from the `amt` package. We annotated the tracks with environmental variables measuring population density (Center for International Earth Science Information Network (CIESIN) Columbia University and CIAT, Centro Internacional de Agricultura Tropical 2005), elevation (U. S. / Japan ASTER Science Team 2009), and landcover class (Defourny et al. 2009) at the end of each simulated and observed step. The original landcover data associated with deciduous, coniferous, or mixed forest were grouped to form a single `forest` variable.

We formed strata by matching each observed step to its 5 associated random steps and fit a conditional logistic regression model with forest, population density, and elevation. We also included step length, log(step length), and cos(turn angle) in the model to update the movement parameters after accounting for habitat selection (Avgar et al. 2016, Fieberg et al. 2021). We then used the `redistribution_kernel` and `simulate_path` functions in the `amt` package (Signer et al. 2023) to simulate fisher movements starting at the same initial location as the 3rd fisher, i.e., the fisher that was not used to parameterize the model. We simulated 19 data sets over the same time interval as the observed data. We matched the pattern of missingness in the observed and simulated trajectories by only including steps with matching timestamps in both the observed and simulated data sets.

We created four different graphical visualizations summarizing the observed and simulated trajectories: 1) a plot of the trajectory overlayed on a map of elevation (Figure 1); 2) a histogram depicting the elevations at the locations visited by the real and simulated animals (Figure 2); 3) a histogram depicting the distribution of turn angles (Figure 3); and 4) a scatterplot depicting the relationship between step lengths and turn angles (Figure 4). The first 3 plots were constructed using functions in the `ggplot2` package (Wickham 2016). The last plot, constructed using the `plot_joint_scat` function in the `cylcop` package (Hodel and Fieberg 2021), is useful for evaluating whether step lengths and turn angles are cross-correlated (Hodel and Fieberg 2022). In addition, we used the `patchwork` package (Pedersen 2020) to stitch together the scatterplots into a single multi-panel plot, the `ggtext` package (Wilke and Wiernik 2022) to annotate this plot, and the `nullabor` package (Buja et al. 2009, Wickham et al. 2010) for helper functions used to create the lineups. A key for identifying the real data in each Figure is provided in the appendix.

**Figure 2:**
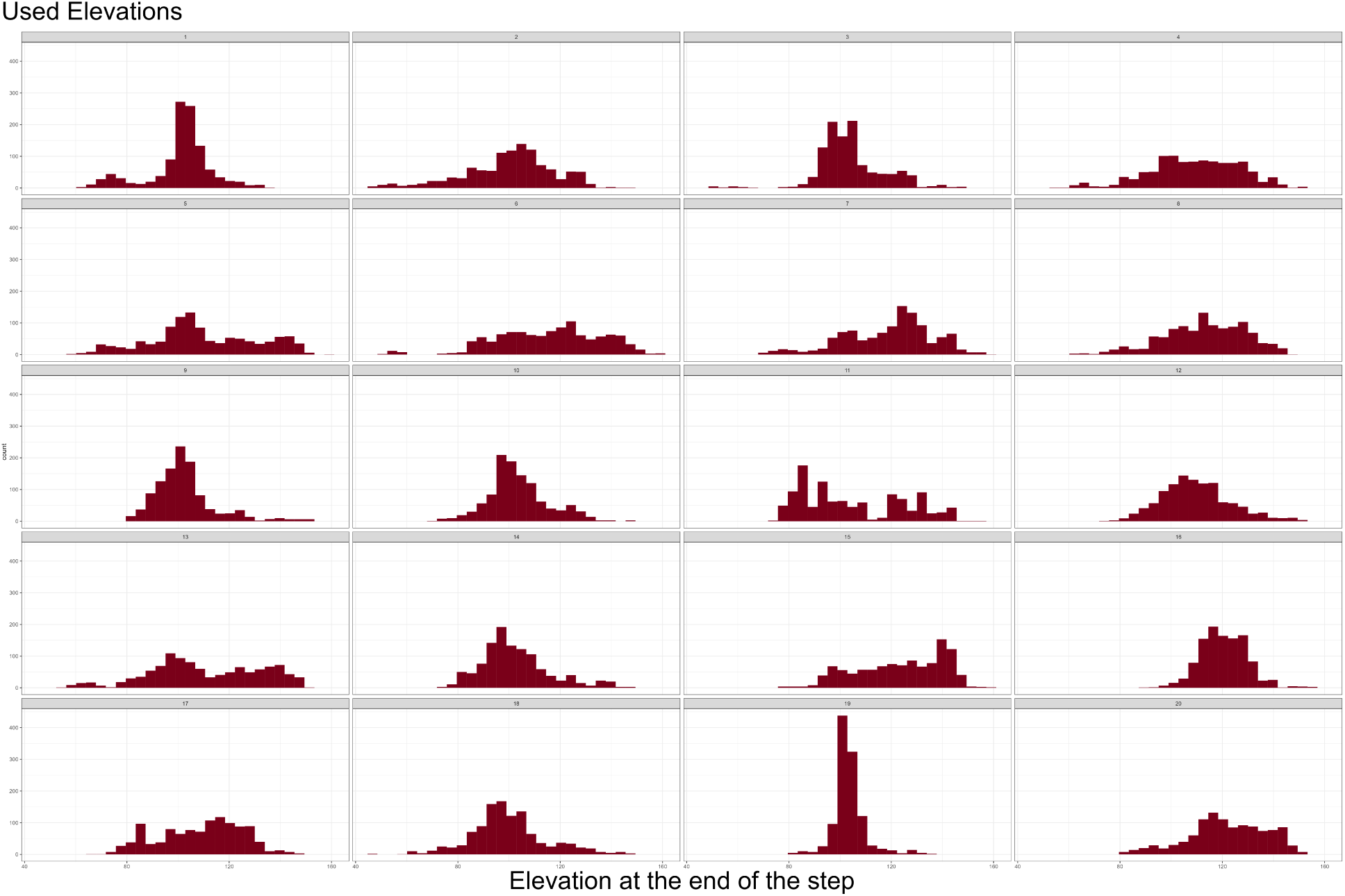
Distribution of elevation measured at the end of each movement step. Nineteen of the panels were generated by simulating observations from an integrated step-selection analysis with fisher data from LaPoint et al. (2013). The other panel contains the distribution of elevations visited by an individual from the same study but whose data were not used to parameterize the model.

**Figure 3:**
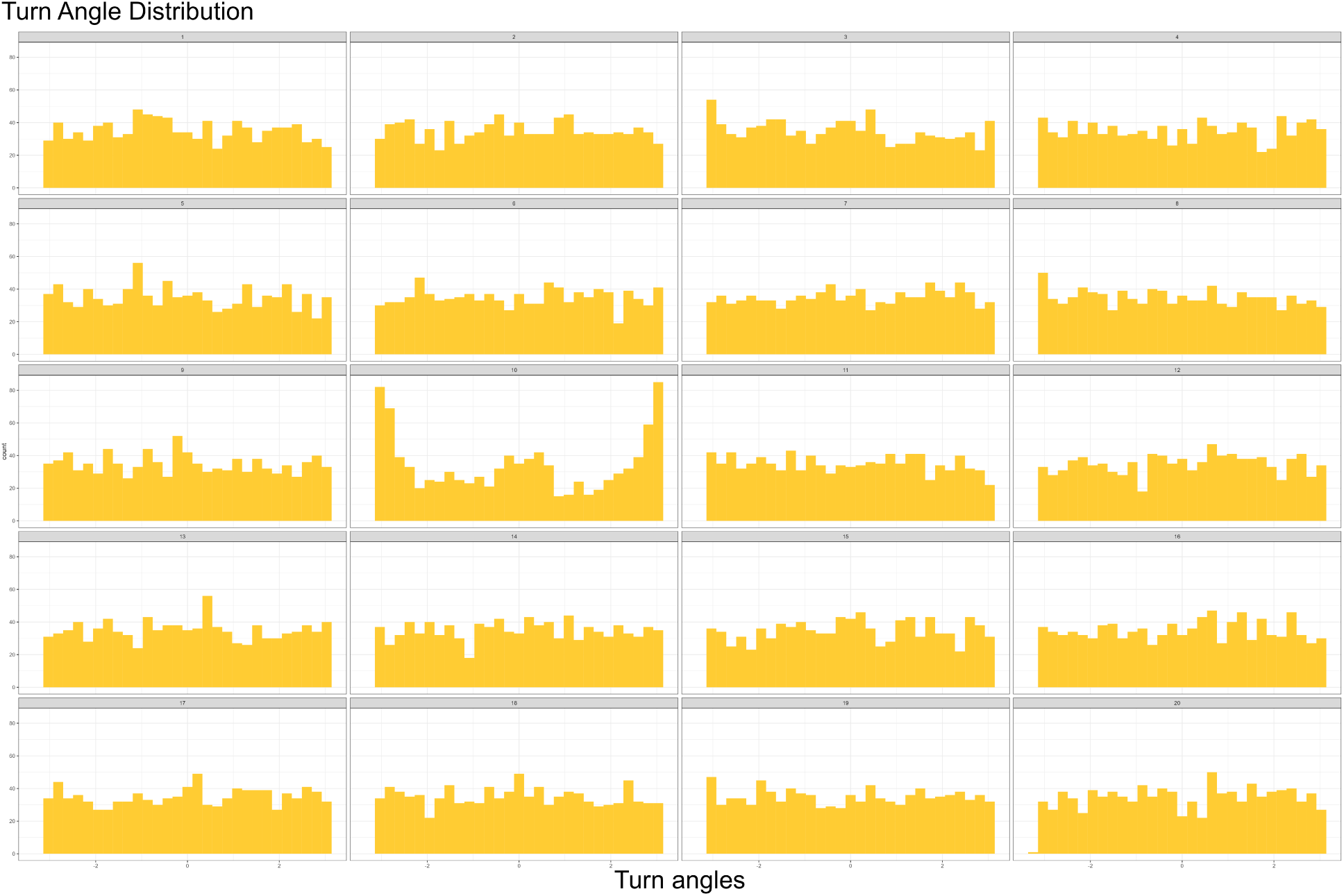
Distribution of turn angles. Nineteen of the panels were generated by simulating observations from an integrated step-selection analysis with fisher data from LaPoint et al. (2013). The other panel contains observed turn angles from movements of an individual from the same study but whose data were not used to parameterize the model.

**Figure 4:**
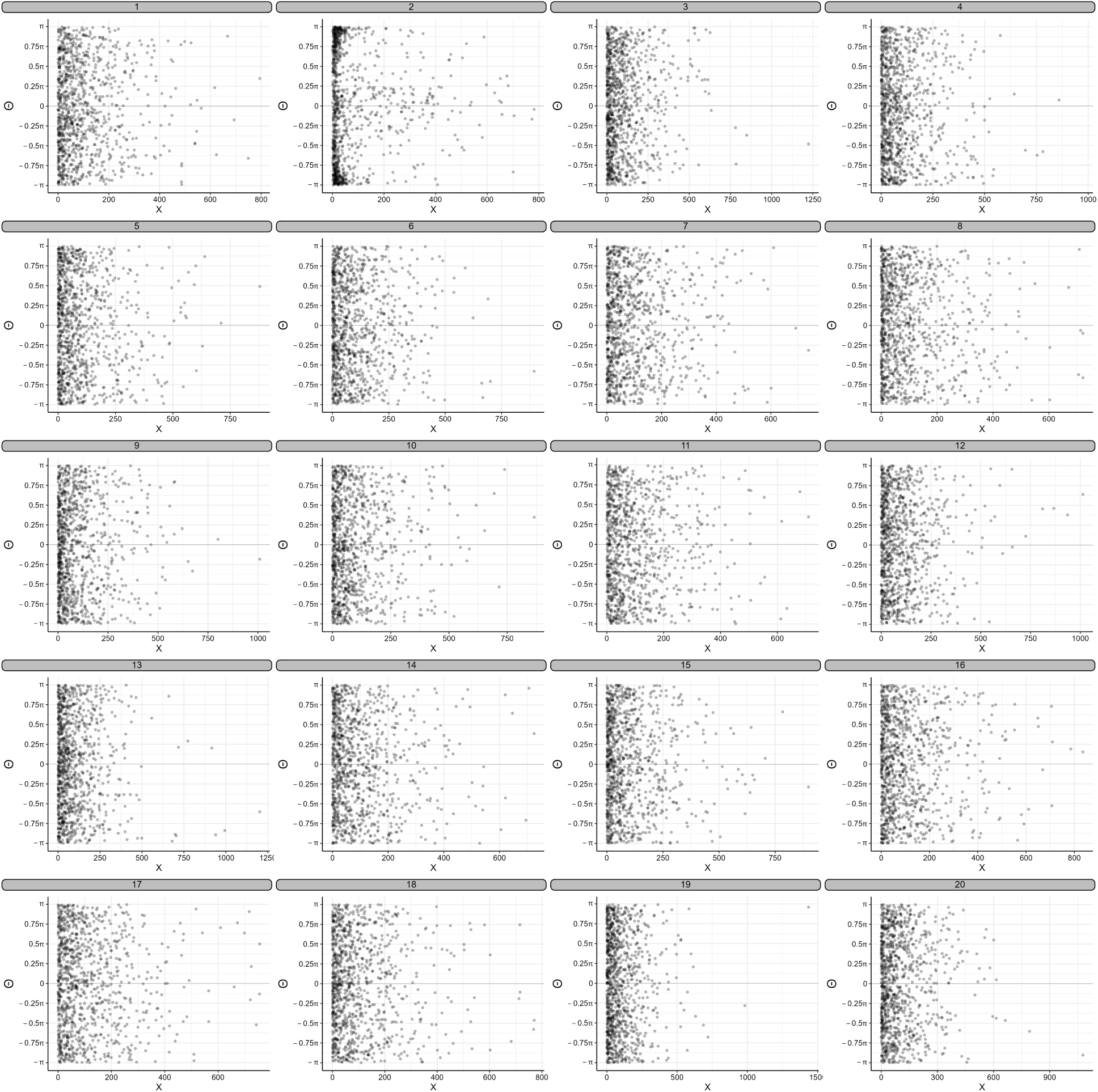
Scatterplot of step lengths (x-axis) and turn angles (y-axis). Nineteen of the panels were generated by simulating observations from an integrated step-selection analysis with fisher data from LaPoint et al. (2013). The other panel contains observed turn angles and step lengths from movements of an individual from the same study but whose data were not used to parameterize the model.

Together, the lineups demonstrate the flexibility of the lineup approach – one can choose *any* graphical summary of the data to test goodness-of-fit. The simulated trajectories provide a quick check on whether the model is capable of capturing various features in the animal’s movements (e.g., whether they are constrained by linear features), and the overall dispersion of the track can also be informative if the *x* and *y* axes are scaled similarly in all panels. In addition, the different simulations can inform us about the range of potential future movements that we might expect to see, assuming the model adequately captures the most important mechanisms driving animal movements. Similar to used-habitat-calibration plots, we can evaluate whether the model predicts the types of locations that are likely to be visited by the animal in the future by plotting the distribution of environmental features at the observed and simulated locations (Figure 2). Lastly, if there are particular features that we hope the model will replicate or assumptions about which we are particularly concerned, then we can construct a custom plot (and test) to address the concern. The observed data are fairly easy to pick out in Figure 3, demonstrating that the von Mises distribution does a poor job at matching the observed turn-angle distribution. Similarly, the observed data in our scatterplot (Figure 4) are fairly conspicuous, which reflects the fact that our ISSA fails to induce a correlation between step lengths and turn angles.

## 4 Hidden Markov Model (HMM) Fit to Fisher Data

We fit a two-state HMM to the same resampled data that we used in the ISSA, but ignored the covariates. We represented each “burst” of steps containing successive locations that were equally-spaced in time as separate individuals and removed “bursts” that contained fewer than 3 locations. We assumed movements were influenced by two behavioral states, which we interpret as representing movements while foraging or travelling. The former state was characterized by less directed movement (i.e., shorter step lengths and larger turn angles) and the latter state by more directed movement (i.e, longer step lengths and shorter turn angles). Conditional on the latent state, we assumed that the distributions of step lengths and turn angles followed Gamma and von Mises distributions, respectively.

To provide an example of how one could easily generate independent lineups for each observer, we fit the same model to three subsets of the data. Each subset included data from a unique combination of two individual fishers, with the third fisher’s data withheld for cross-validation. We used the `simData` function in the `momentuHMM` package (McClintock and Michelot 2018) to simulate trajectories for the individual fisher that was not used to parameterize the model in the given iteration. The subsequent lineups (Supplemental Figures S1, S2, and S3) could then be shown to three independent observers and the p-value calculated as outlined in Section 2.

By constructing the same set of plots for both HMMs and ISSAs, we can compare their relative goodness-of-fit. For example, in Figure 5, we contrast simulated data from our HMM and our ISSA to evaluate their ability to capture the observed cross-correlation between fisher step lengths and turn angles. Rather than use a scatterplot as in Figure 4, this time we consider circular boxplots constructed using the `plot_joint_box` function in the `cylcop` package (Hodel and Fieberg 2021). The colored boxes capture the middle 50% of the distribution of turn angles for different quantiles of step length, with the innermost boxplots corresponding to the smallest step lengths. For the HMM (left two columns), the distribution of turn angles is centered on 0 for the outer rings corresponding to the largest step lengths and centered on *π* for the innermost rings corresponding to the smallest step lengths. By contrast, the boxplots associated with the ISSA (right two columns) show no consistent pattern with the different quantiles of step lengths. This example highlights that the latent states of the HMM induce cross-correlation between step lengths and turn angles that we also see in the observed data, whereas the ISSA is not able to capture this feature of the data. It also serves to demonstrate that multiple visualizations can be constructed to graphically assess the same assumption (e.g., Figure 4 or Figure 5), and users are free to choose the visualization that they prefer.

**Figure 5:**
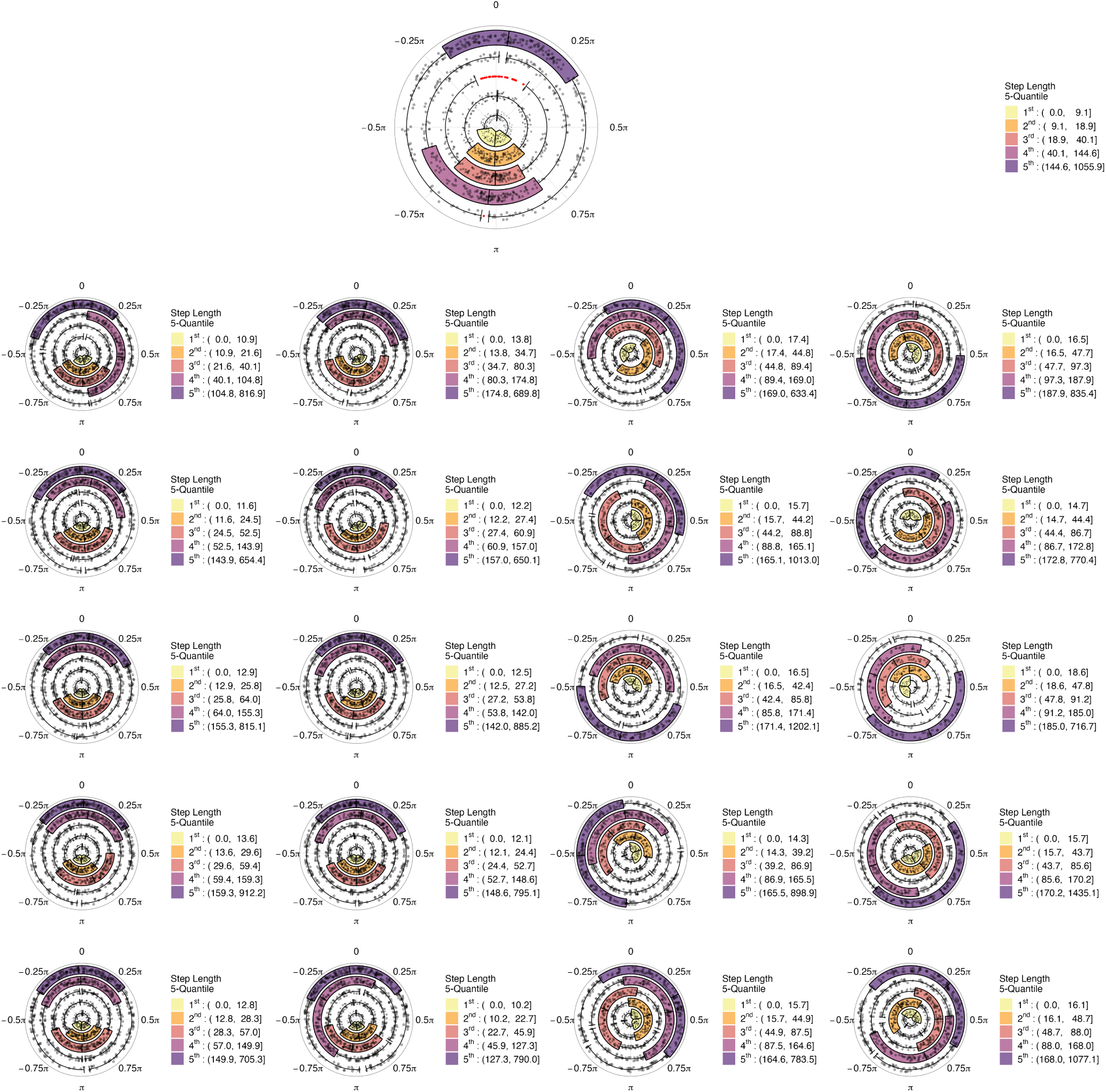
Distribution of turn angles for different quantiles of step lengths. The top panel is the observed movement data of the individual not used to parameterize the model. The left two columns contain plots generated by simulating observations from the the two-state HMM. The right two columns contain plots generated by simulating observations from from the ISSA model. Both models were fit with the same fisher data from LaPoint et al. (2013)

## 5 Discussion

We have shown how simulations from animal movement models can be used to form lineups for evaluating model goodness-of-fit. We focused on integrated step-selection analyses and hidden Markov models due to their widespread use and utility. ISSAs result in a *spatially-explicit*, individual-based model of animal movement capable of capturing a wide range of mechanisms, including individual responses to local environmental features such as roads and seismic lines (Prokopenko et al. 2017, Scrafford et al. 2018, Dickie et al. 2020), interactions with conspecifics (Schlägel et al. 2019), and familiarity with previously visited locations and migration routes (Oliveira-Santos et al. 2016, Merkle et al. 2019, Kim et al. 2023). As such, ISSAs offer a powerful approach for parameterizing individual-based models of animal movement that can be simulated to model connectivity (Hofmann et al. 2023) or to predict how animals will respond to land use or climate change (Signer et al. 2017, Potts et al. 2022). HMMs, although not spatially explicit, offer a computationally efficient approach for inferring otherwise unobservable animal behavior from telemetry and environmental data (Langrock et al. 2012). Applications of HMMs range from cataloging harbor seal activity budgets (McClintock et al. 2013) to reconstructing bird trajectories based on pressure and wind data (Nussbaumer et al. 2023).

Despite the widespread use of ISSAs and HMMs in ecology, there are relatively few methods available for assessing their goodness-of-fit; we hope that lineups can help fill that gap. An important consideration when evaluating models, particularly those used in conservation and management, is whether they are capable of generating accurate predictions across space or time, referred to as *model transferability* (Yates et al. 2018, Helmstetter et al. 2021, Rousseau and Betts 2022). By treating individuals as independent sample units for cross-validation, we can test how well models fit to individuals from a restricted subset of environments perform when evaluated using individuals living elsewhere. Thus, lineups, and particularly those that visualize the types of environments visited by simulated and observed animals (e.g., Figure 2), seem particularly appealing for evaluating model transferability (Fieberg et al. 2018).

In our application of lineups to the fisher data, we found that the ISSA was capable of generating simulated trajectories that looked like the real trajectory for the “out-of-sample” fisher (Figure 1). However, some of the simulated trajectories looked quite different from the observed trajectory, and it failed to reproduce the observed peak at *±π* in the distribution of turn angles (Figure 3) or the correlation between observed step lengths and turn angles (Figures 4, 5). The HMM was able to better capture these features of the data by modeling movements as a function of a latent behavioral state taking on one of two discrete values. These states allow for a mixture of movement types (long and directed vs. short and tortuous), which in turn induces correlation between step lengths and turn angles (Hodel and Fieberg 2022). Building on these two approaches, one might next consider fitting a state-switching step-selection function (Klappstein et al. 2023, Pohle et al. 2023), though the software for fitting and simulating from this class of models is less well developed.

We demonstrated, via the HMM application, how lineups can be replicated and shown to multiple users. In the past, some researchers have used Amazon Mechanical Turk for this purpose (Hofmann et al. 2012, Majumder et al. 2014), though we suggest participatory science platforms, such as Zooinverse (Simpson et al. 2014), offer an appealing alternative. If the same lineup is shown repeatedly to different observers, then a binomial sampling distribution will no longer be appropriate for calculating p-values, but model-based inferential frameworks are available (VanderPlas et al. 2021).

Although we have focused on applications of the lineup protocol for goodness-of-fit testing of animal movement models, the approach can be used much more broadly (see e.g., applications in Buja et al. 2009, Wickham et al. 2010). With sufficient observers, lineups have been shown to be competitive with traditional methods for testing hypotheses with linear models (i.e., they have been shown to have similar statistical power) (Majumder et al. 2011, 2013). The advantage of the lineup approach, however, is in its flexibility - i.e., *any* visual summary of the data can be turned into a test statistic. This feature also makes the lineup protocol interesting from a statistical education standpoint, since it offers a unique application of standard null hypothesis testing. Lastly, we see lineups as a valuable tool that can be used in an iterative approach to model building (sensu Potts et al. 2022), whereby features of the data that are poorly replicated can serve to motivate changes in model structure.

## Acknowledgements

JF was supported by National Aeronautics and Space Administration award 80NSSC21K1182 and also received partial salary support from the Minnesota Agricultural Experimental Station.

## Author Contributions

JF conceived of the study, led the integrated step-selection analysis, and drafted the first version of the paper. SF analyzed the fisher data using hidden Markov Models and prepared the vignettes for both applications. JS consulted on the simulation approach. All authors discussed, edited, read, and approved the final manuscript.

## Appendix

This appendix includes additional lineups from the Hidden Markov Model applied to the fisher data and “solutions” to help identify the real data panel in each lineup presented in the paper and in this appendix.

**Figure 6:**
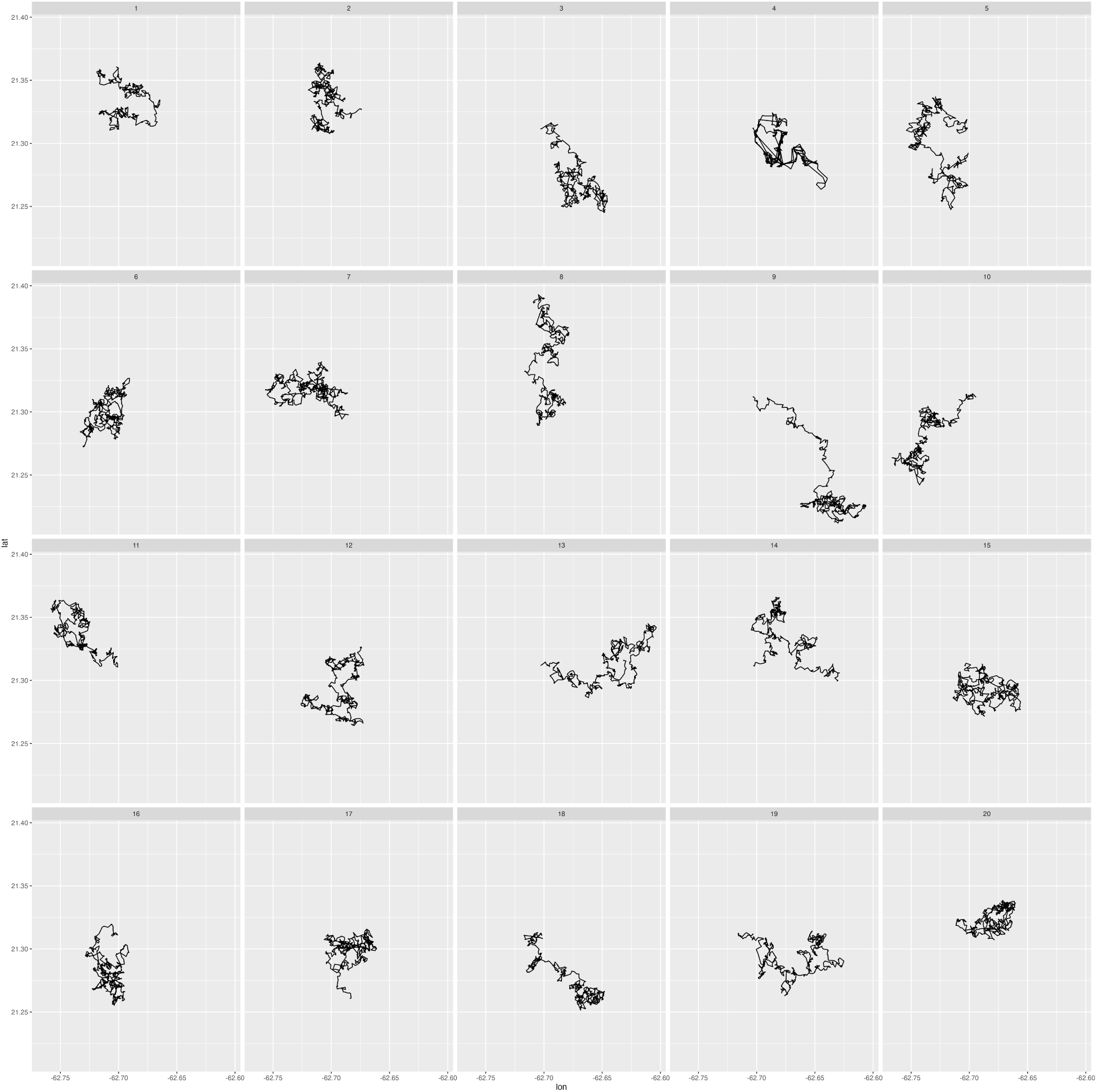
Simulated trajectories. Nineteen of the panels were generated by simulating observations from a 2-state hidden Markov model fit with fisher data from LaPoint et al. (2013). The other panel contains an observed trajectory for the individual from the same study but from data that were not used to parameterize the model.

**Figure 7:**
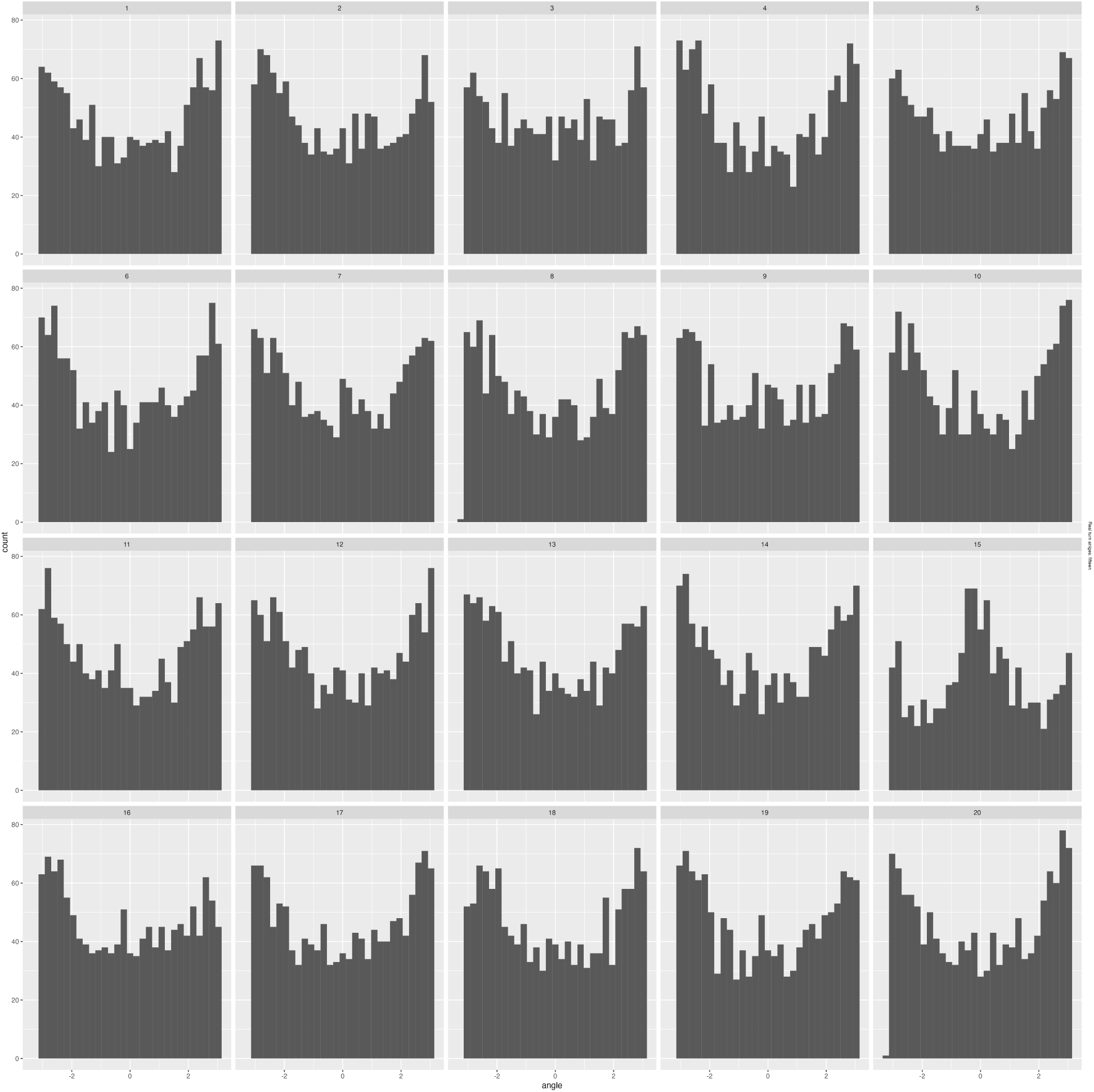
Distribution of turn angles. Nineteen of the panels were generated by simulating observations from a 2-state hidden Markov model fit with fisher data from individuals 1072 and 1078 from LaPoint et al. (2013). The other panel contains observed turn angles from movements of individual 1016.

**Figure 8:**
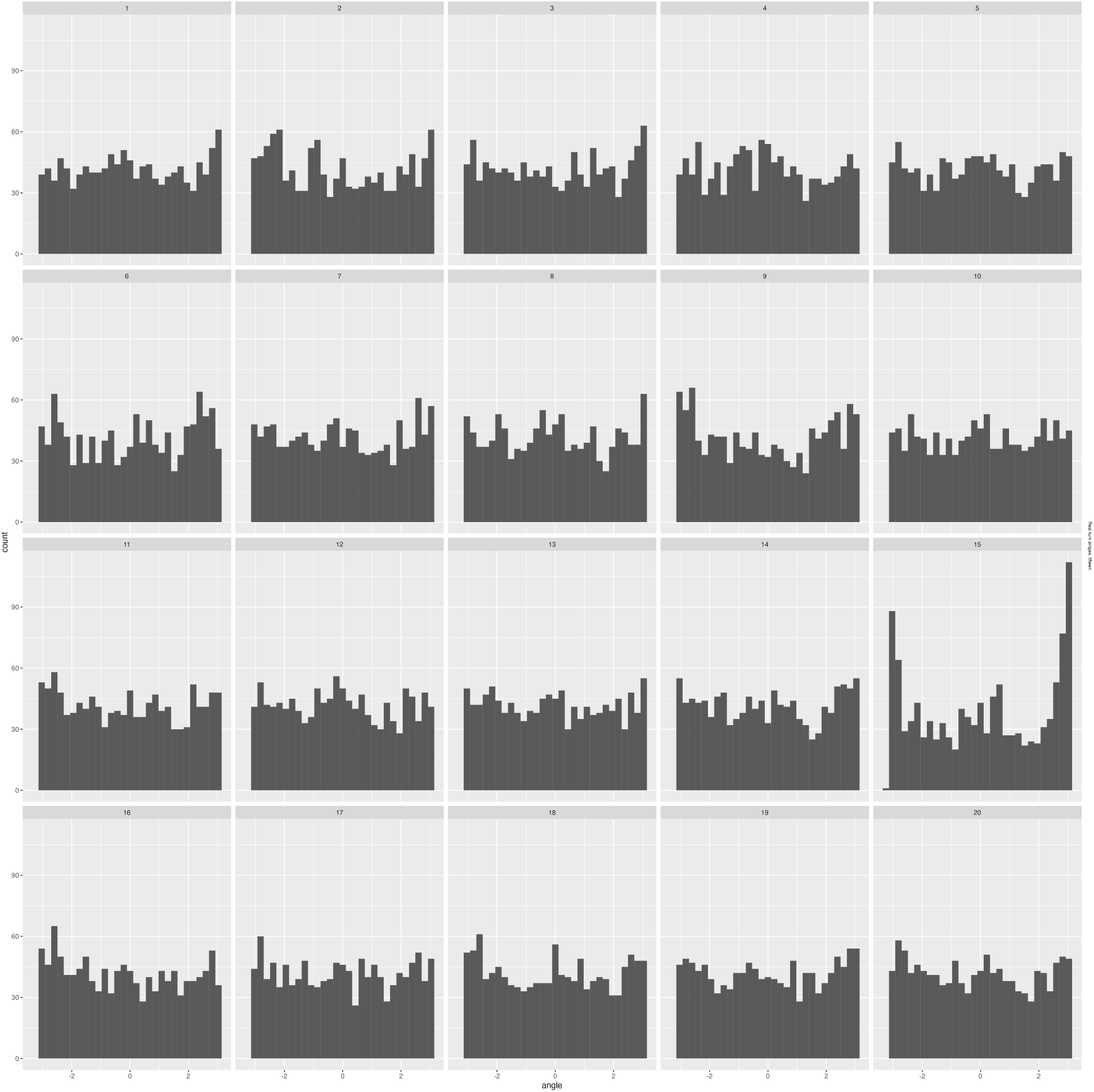
Distribution of turn angles. Nineteen of the panels were generated by simulating observations from a 2-state hidden Markov model fit with fisher data from individuals 1016 and 1078 from LaPoint et al. (2013). The other panel contains observed turn angles from movements of individual 1072.

**Figure 9:**
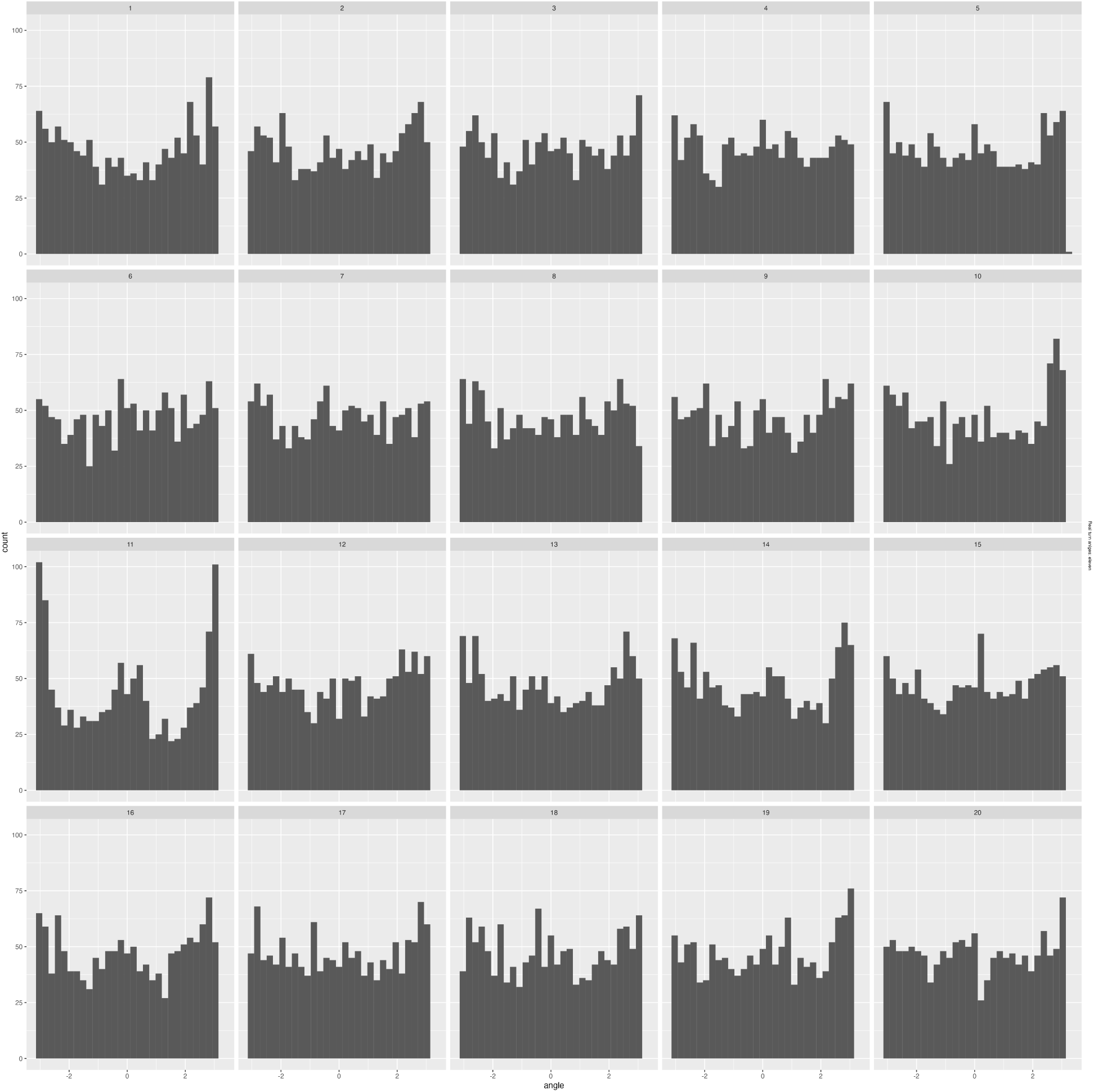
Distribution of turn angles. Nineteen of the panels were generated by simulating observations from a 2-state hidden Markov model fit with fisher data from individuals 1016 and 1072 from LaPoint et al. (2013). The other panel contains observed turn angles from movements of individual 1078.

### Solutions: Will the Read Data Please Stand Up?

Position of observed data in figures:

- Figure 1: the real trajectory is given by panel 6
- Figure 2: the distribution of elevation at the observed locations is depicted in panel 11
- Figure 3: the distribution of observed turn angles is depicted in panel 10
- Figure 4: the distribution of observed step lengths and turn angles is depicted in panel 2
- Figure S1: the real trajectory is given by panel 4
- Figure S2: the distribution of observed turn angles is depicted in panel 15
- Figure S3: the distribution of observed turn angles is depicted in panel 15
- Figure S4: the distribution of observed turn angles is depicted in panel 15

### Using The Lineups Protocol for Exploratory Goodness-of-Fit Evaluation of ISSAs

09-22-2023

#### Document Preamble

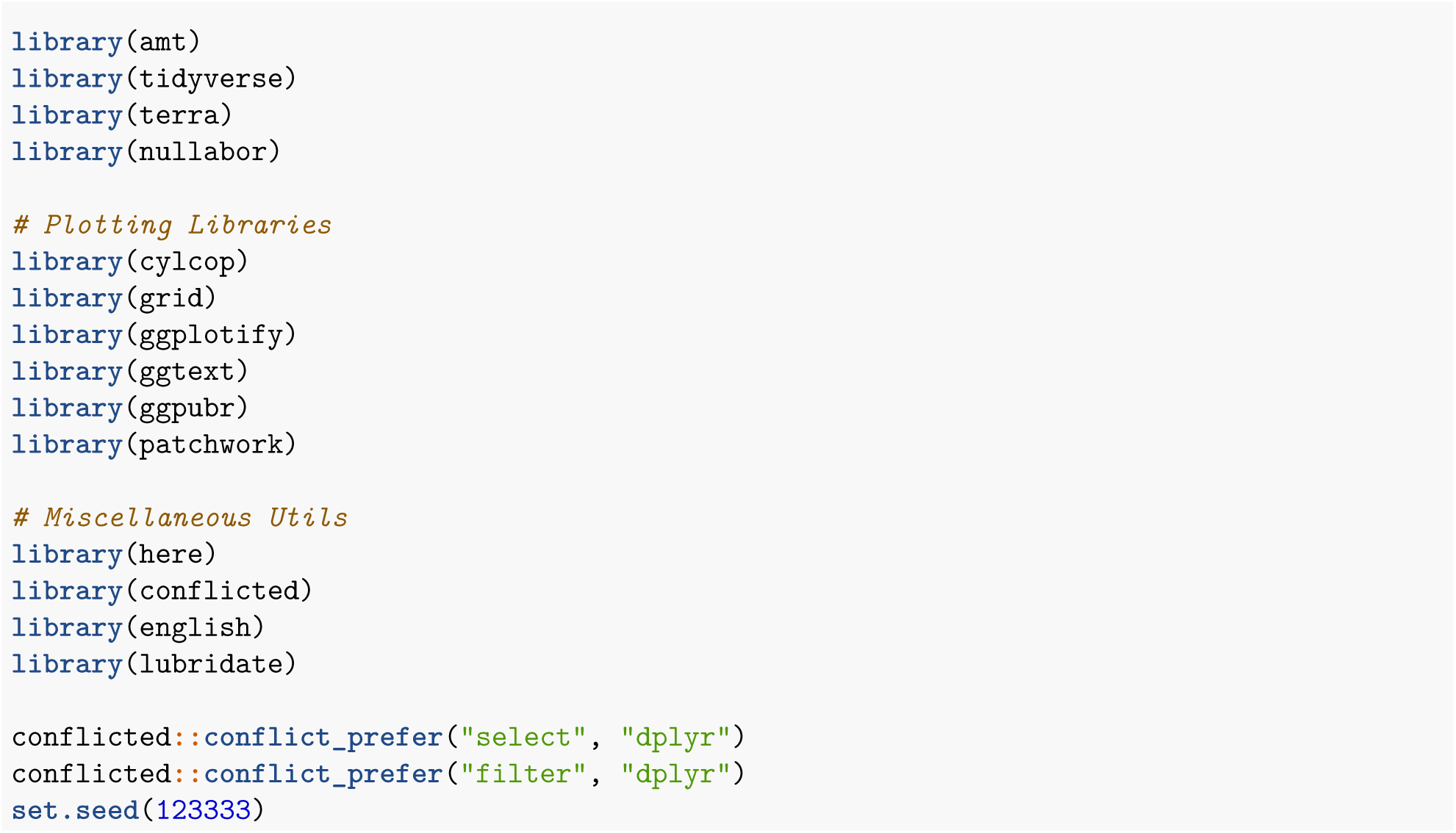

#### Intro

The purpose of this vignette is to demonstrate how to use the lineup protocol in tandem with Integrated Step Selection Analyses (ISSAs).

#### Training and Testing Data

In order to demonstrate a simplistic way to separate our total data into training and test data, we have written the following function, get_data, which takes a given fisher ID whose data will be used to test, the remaining data will be used to parameterize the model. As we show in the HMM vignette (not included here), this function can be used to iteratively fit the same model to different subsets of the data. The following function does three primary things: 1) loads the covariates landcover, population density, and elevation from the amt package, 2) generates 5 random points for each observed point, assuming a gamma distribution for step lengths and a von Mises for turn angles, and 3) divides the data into training_data (what will be used to parameterize the model) and test_data (which will be used to compare model-simulated data and observed data).

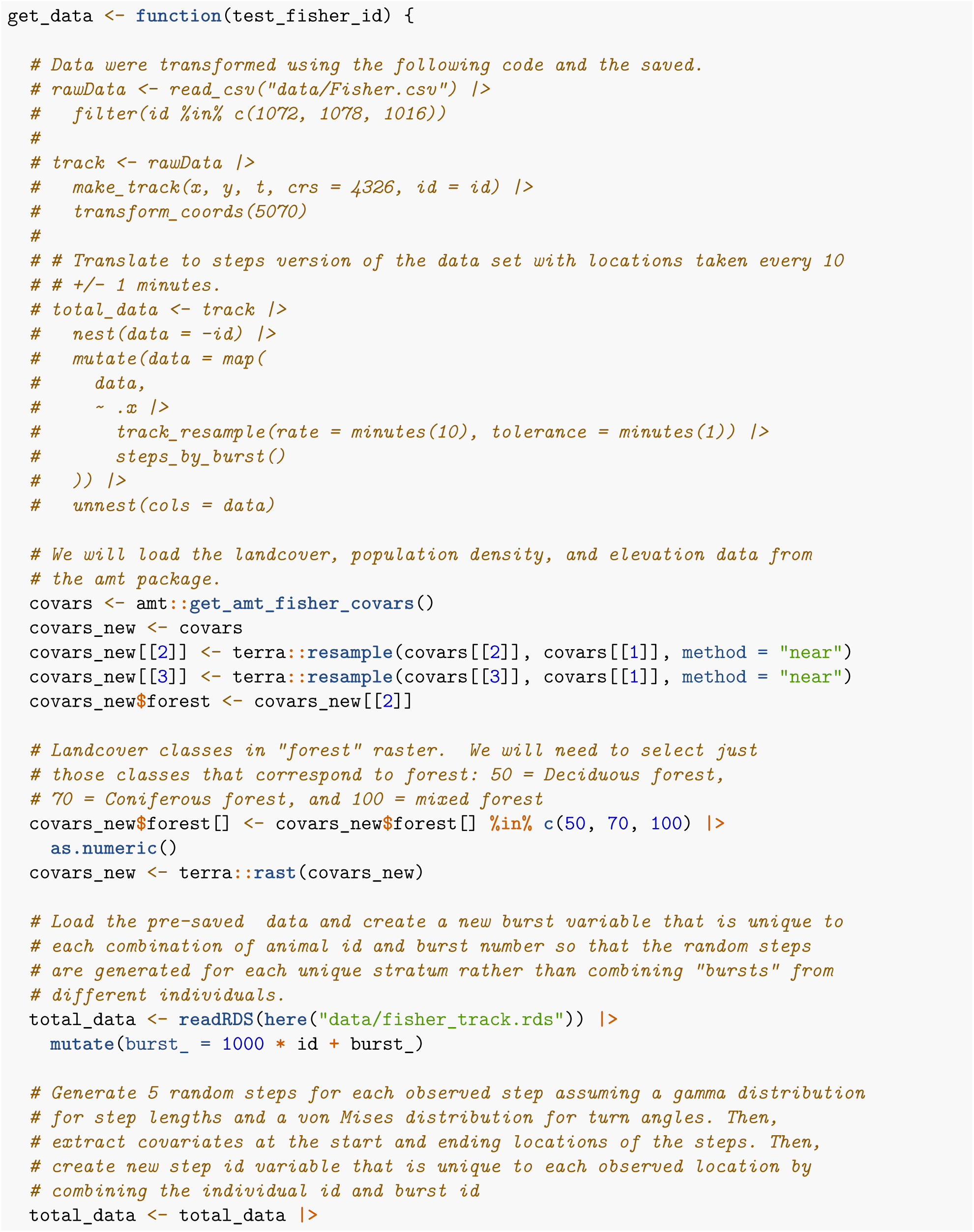
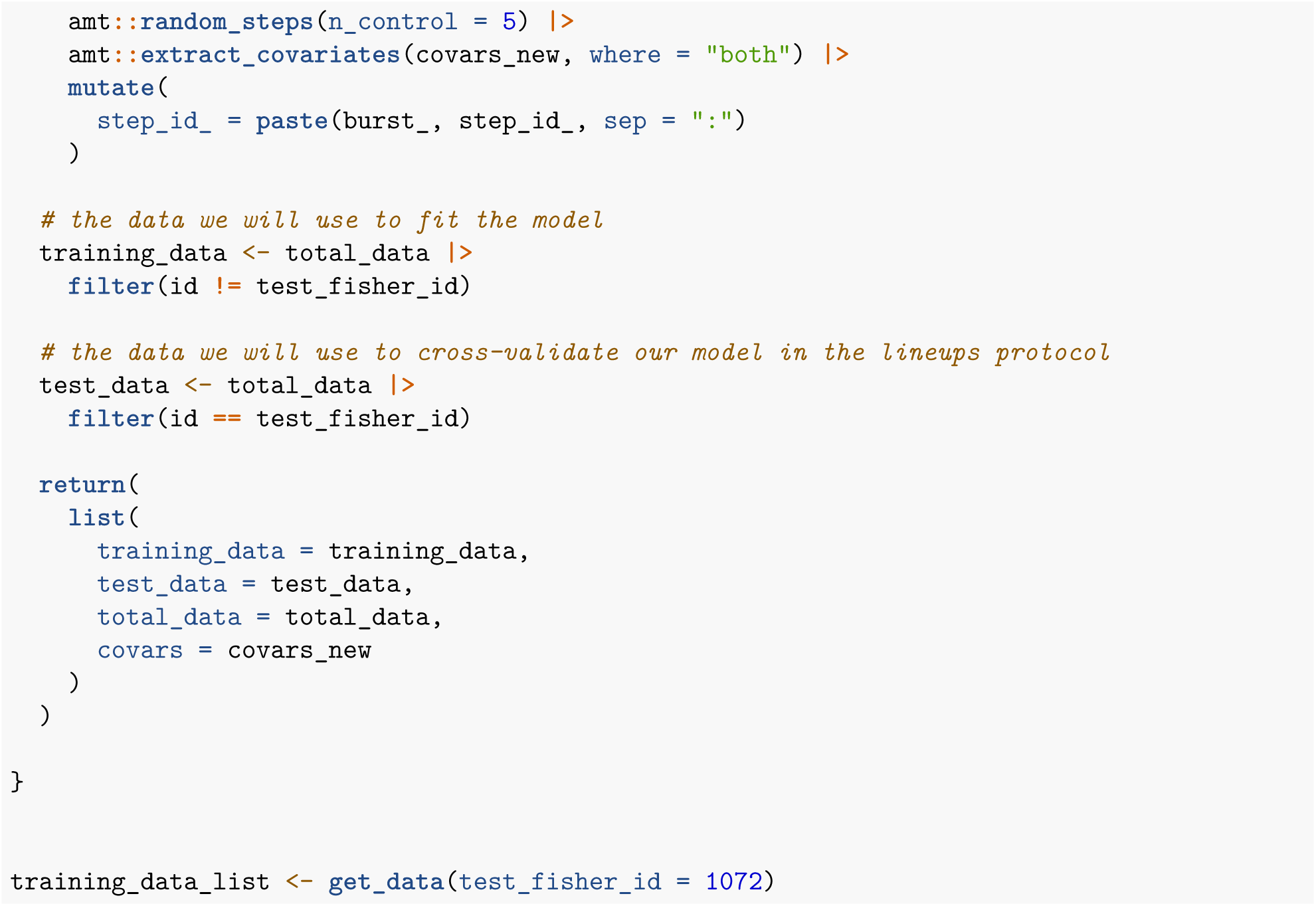

#### Fitting an ISSA

Now that we have our data, let’s actually fit the model using our function fit_model and take a look at the model summary.

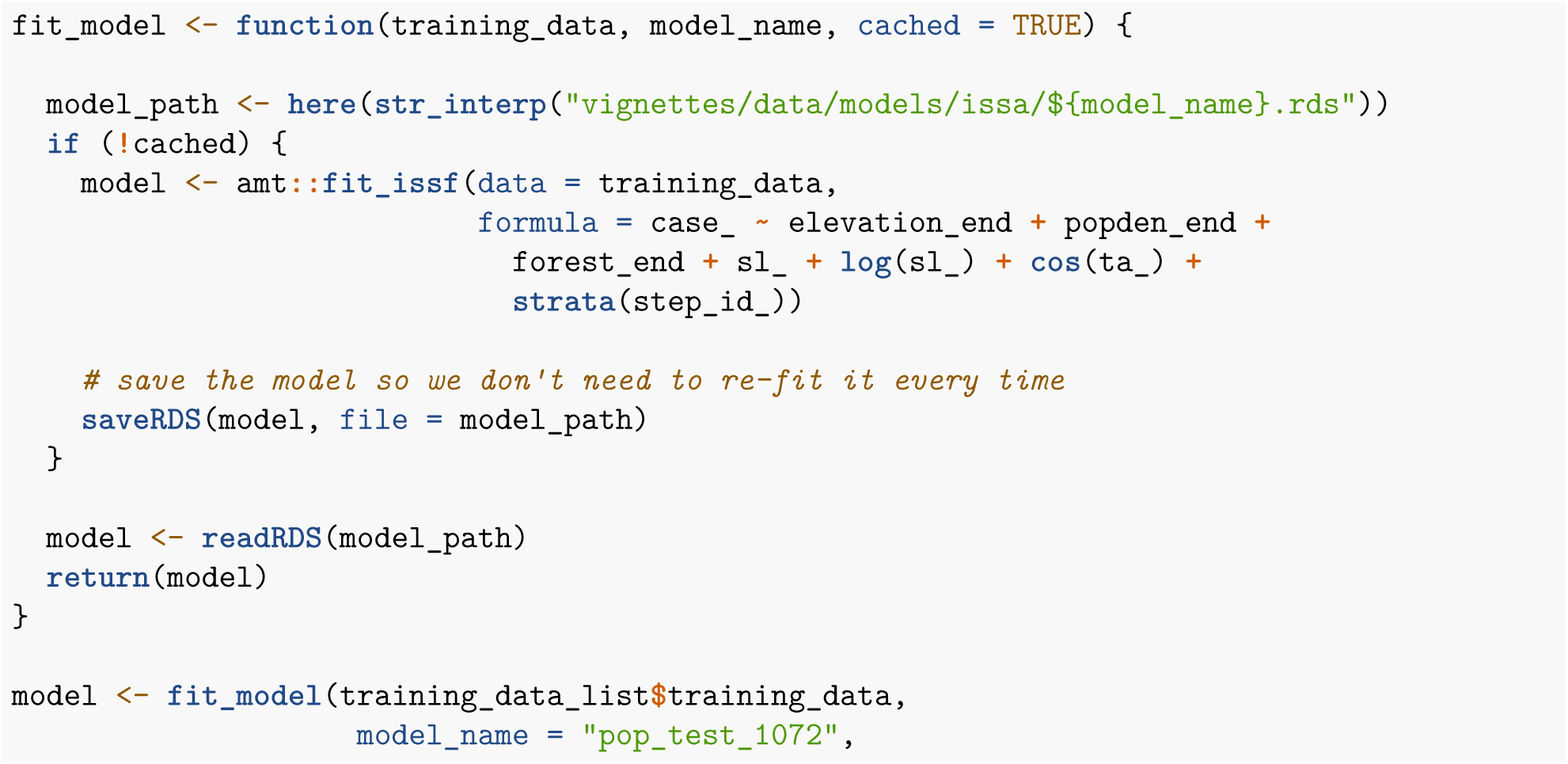
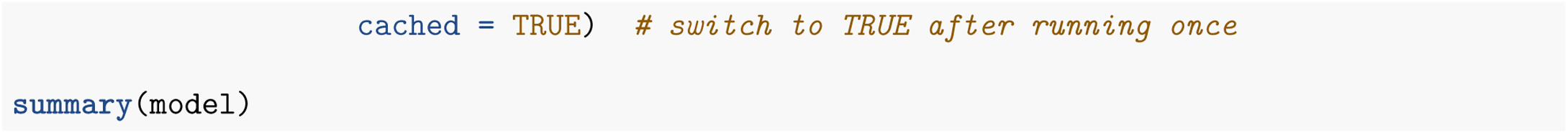

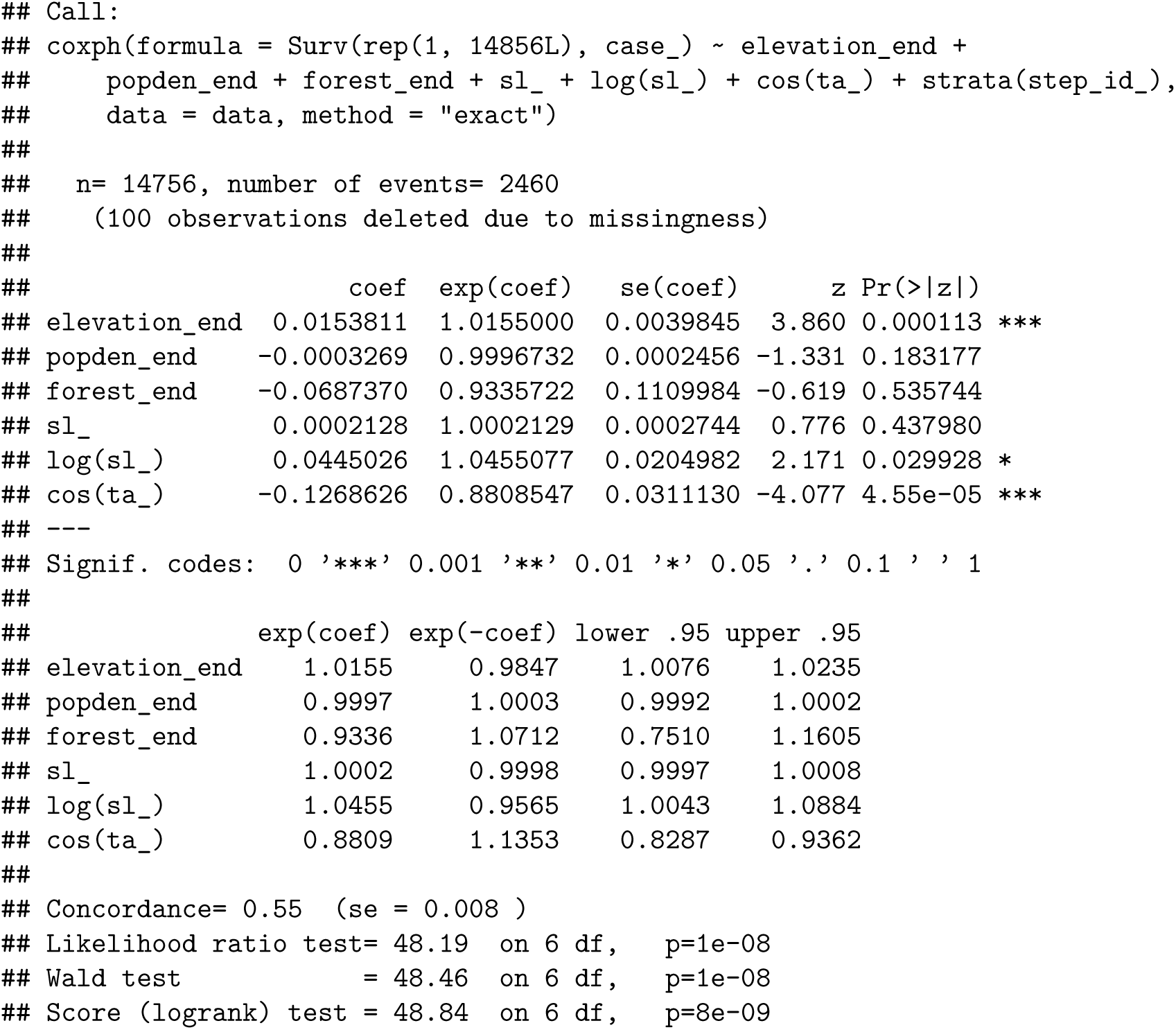

#### Generating Lineup Protocol Artifacts

For the sake of this vignette we will focus on creating used-elevations histograms and step length & turn angle correlation plots. But remember! The lineup protocol can accommodate any visualization of the data.

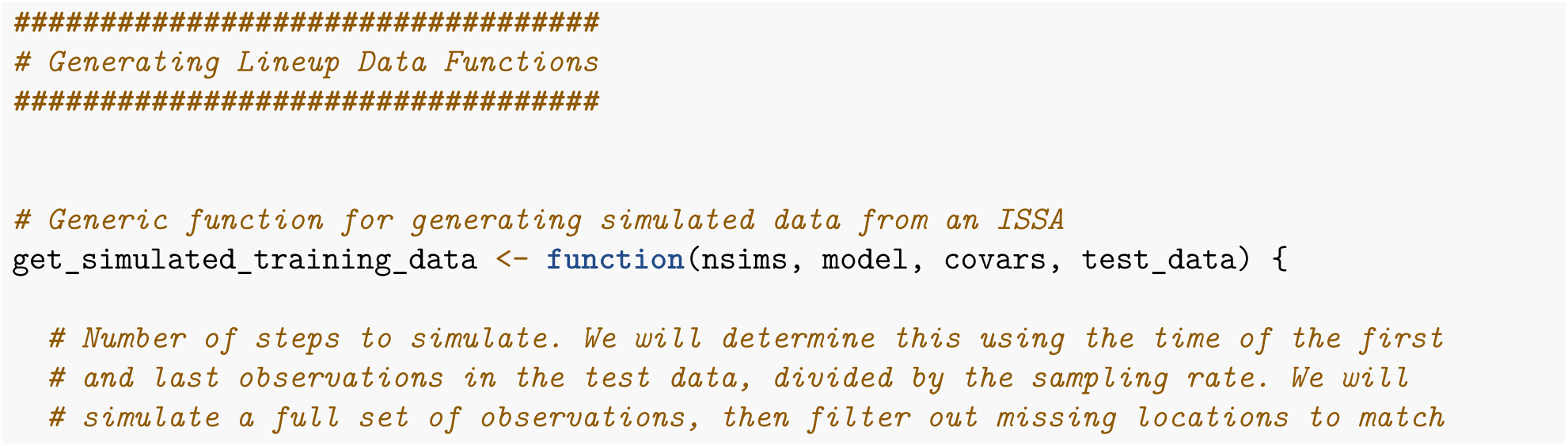
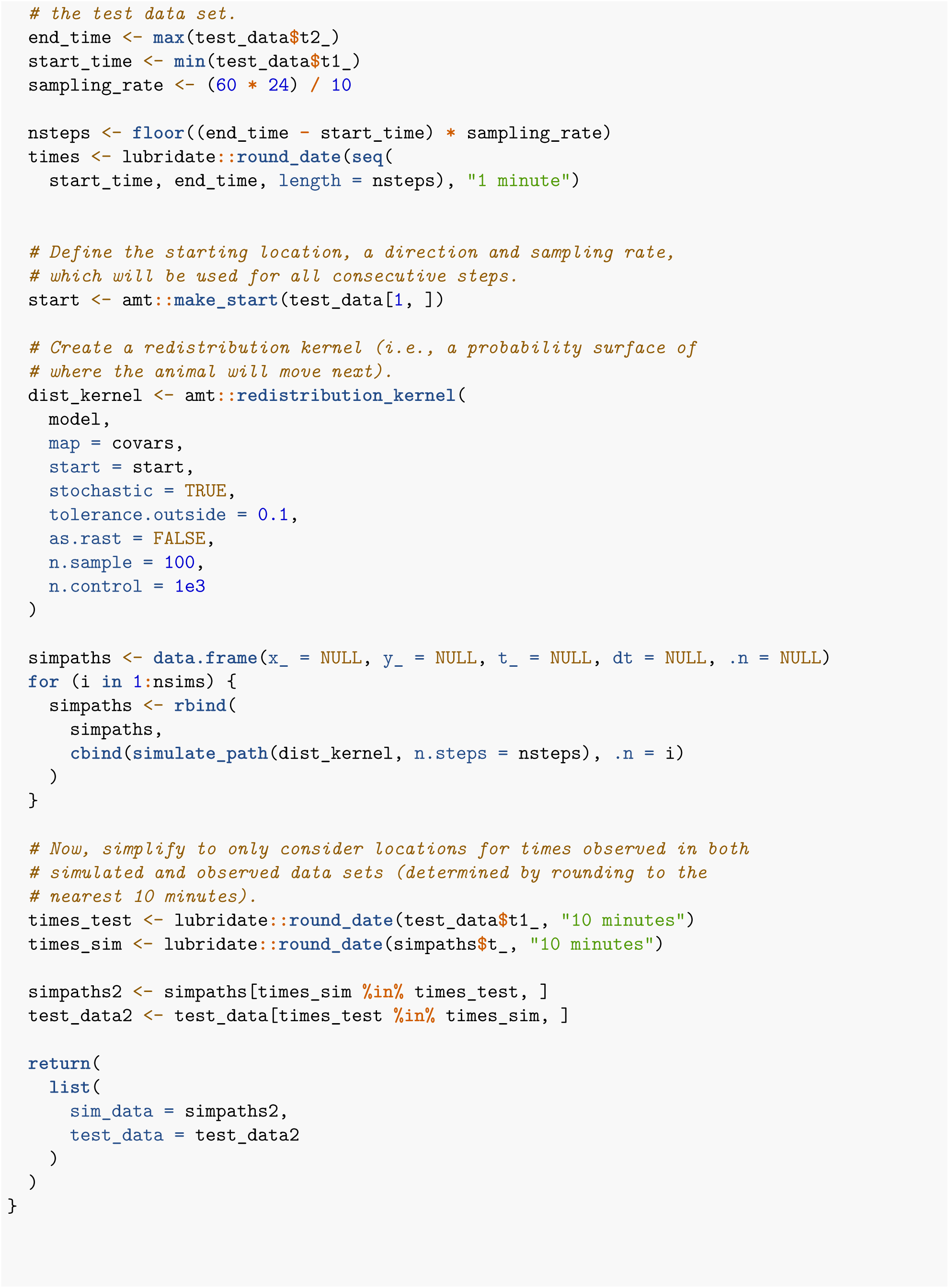
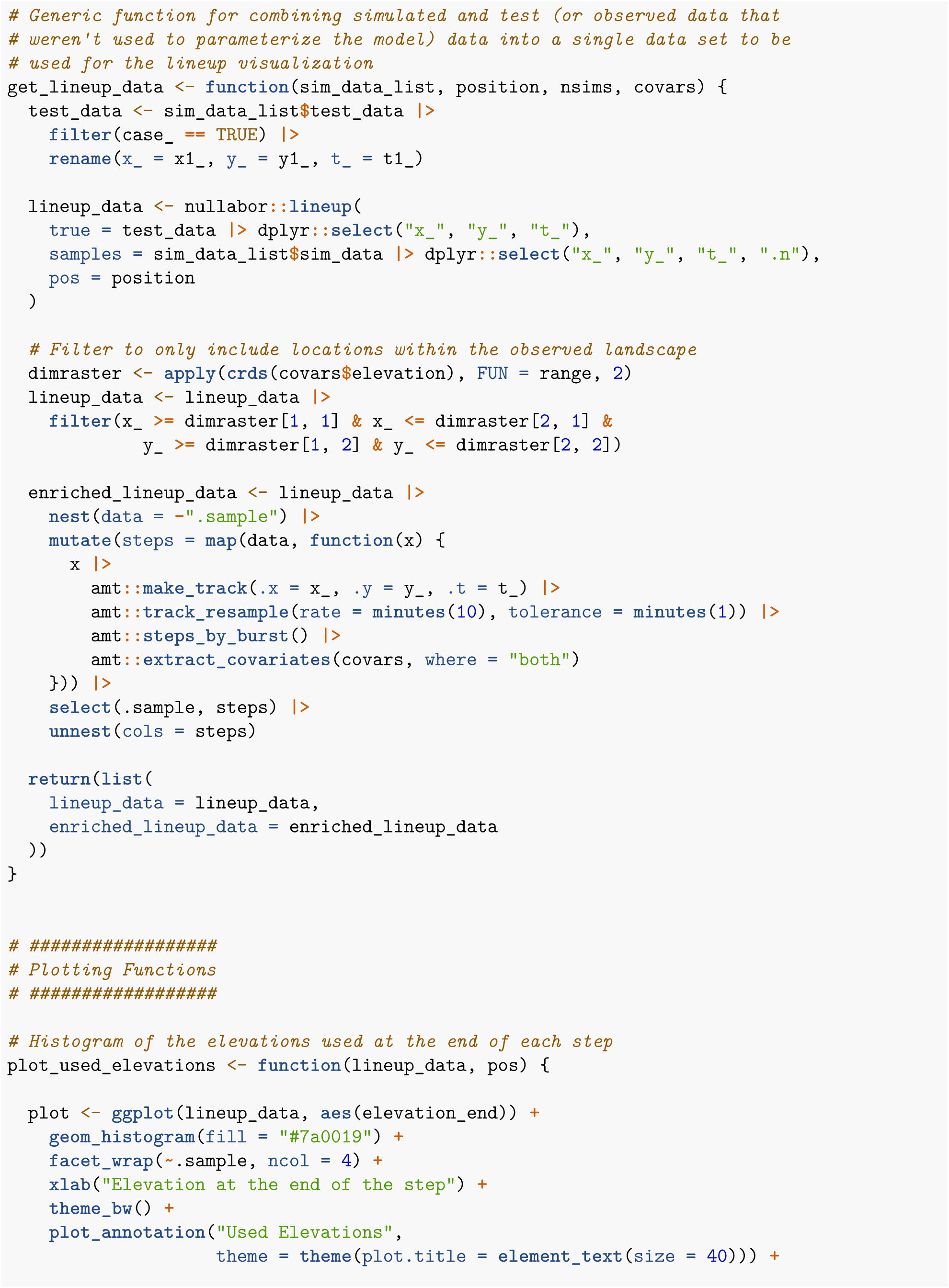
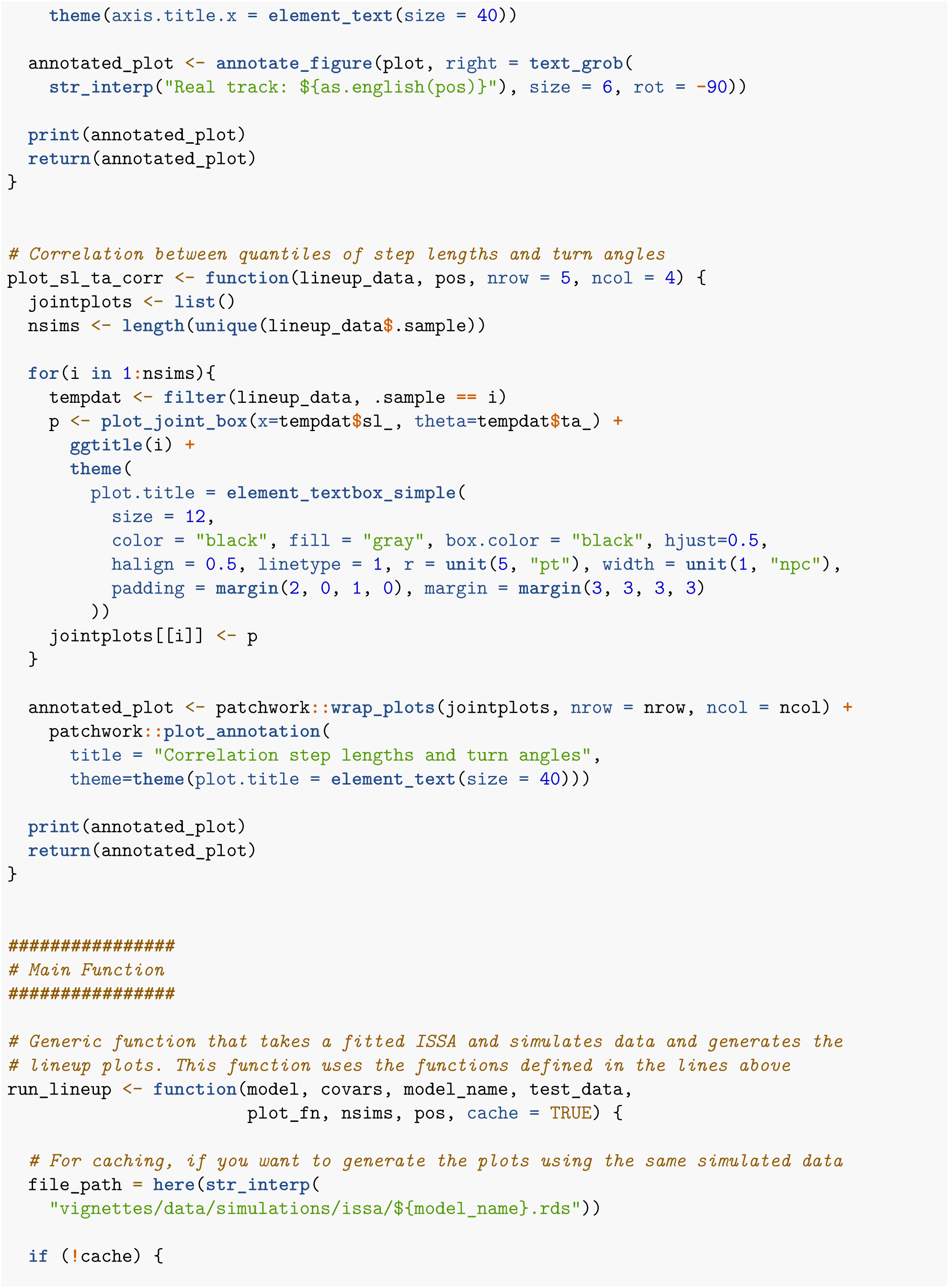
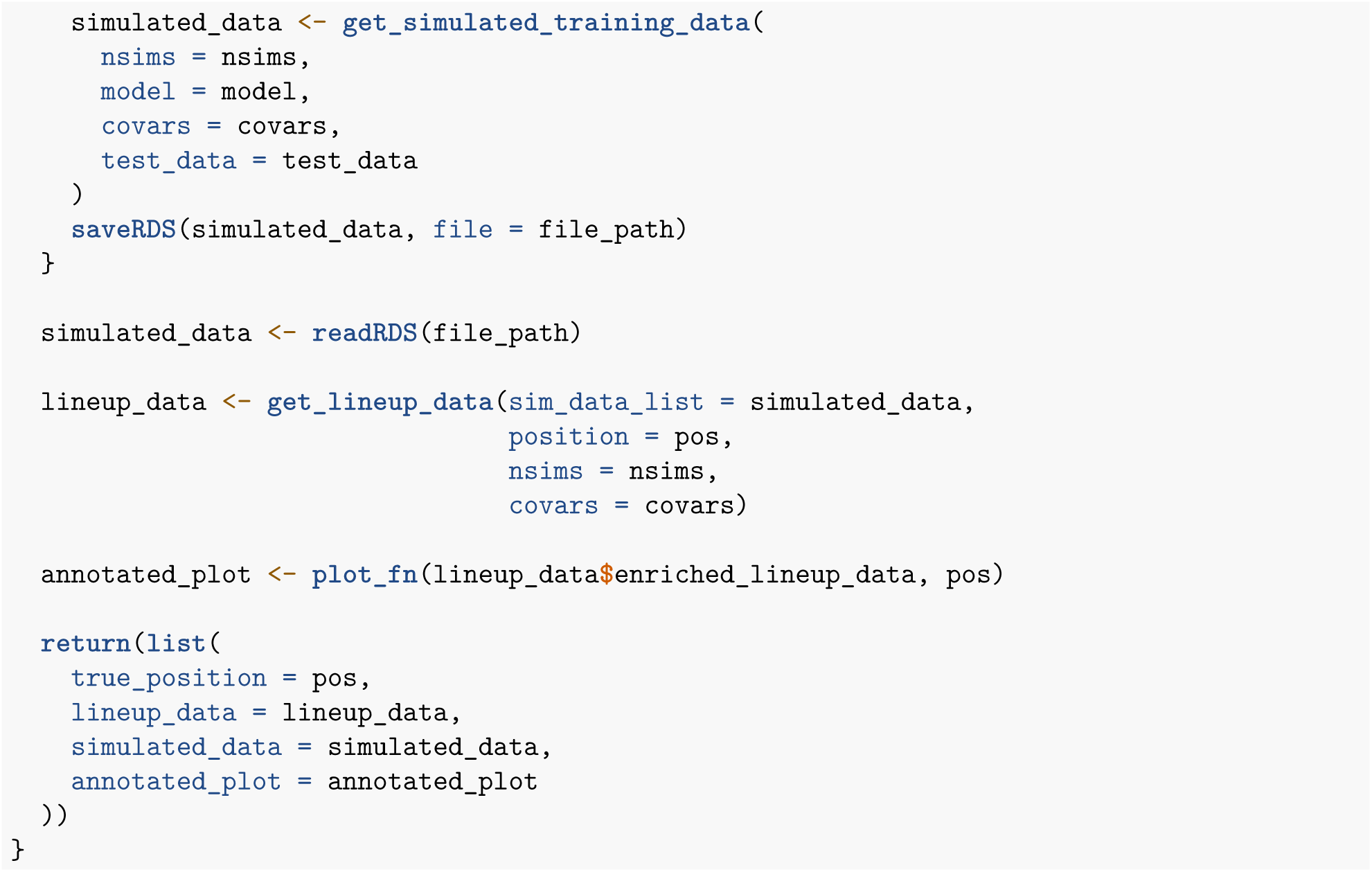

#### Run The Lineup

##### Used Elevations

Now that we’ve defined the methods that will let us generate the lineups in one fell swoop, let’s actually execute the code. First we’re going to take a look at a lineup for used elevations.

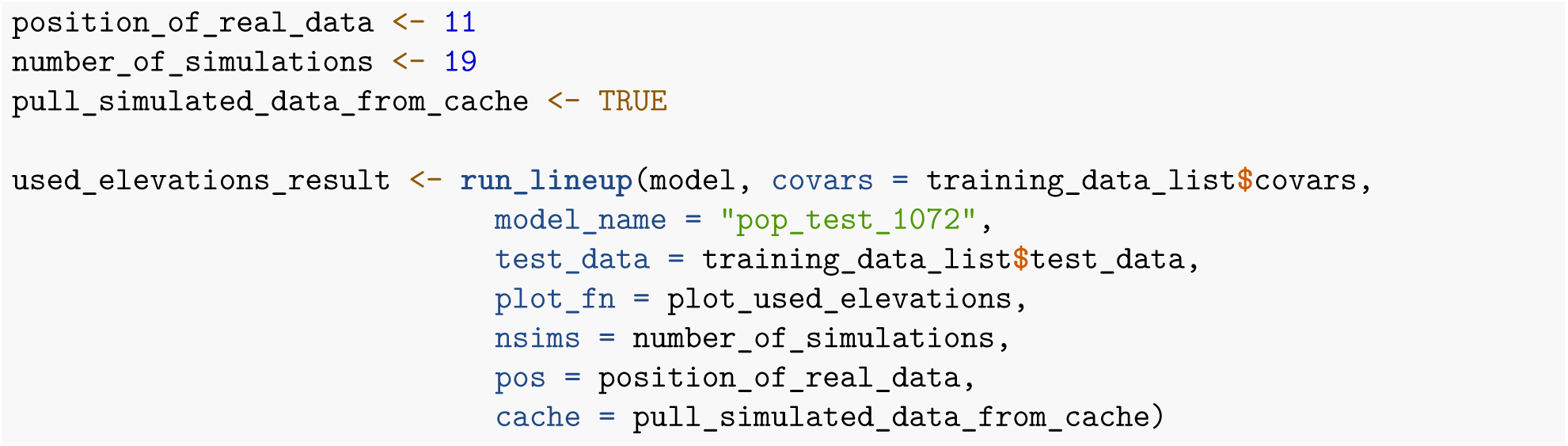

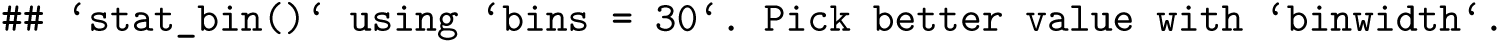

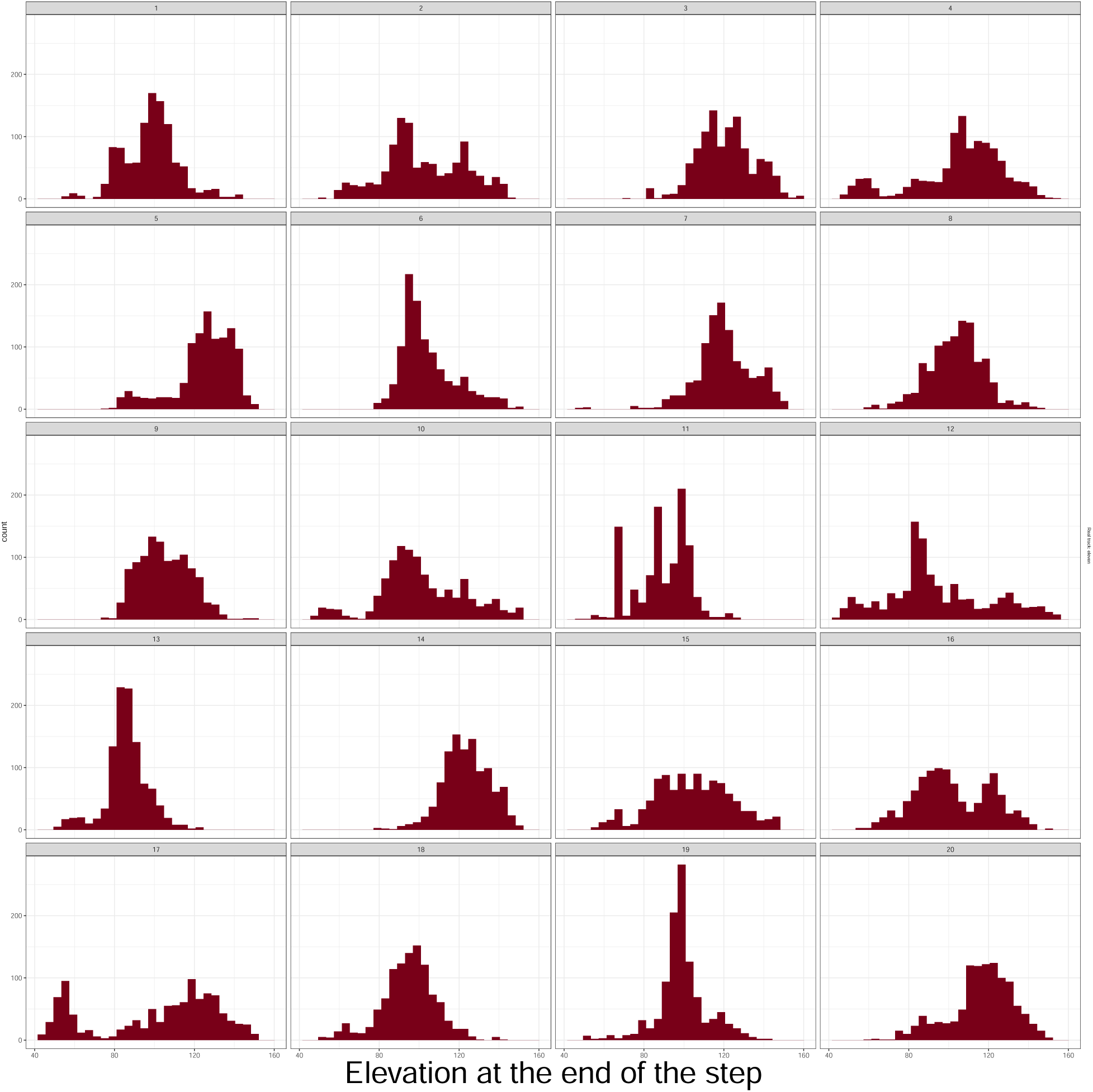

#### Step Length & Turn Angles Correlation

Now let’s look at a lineup for step length and turn angle correlation. In observed data, these variables are auto-correlated, but, as you can see, our ISSA does a poor job of preserving this relationship.

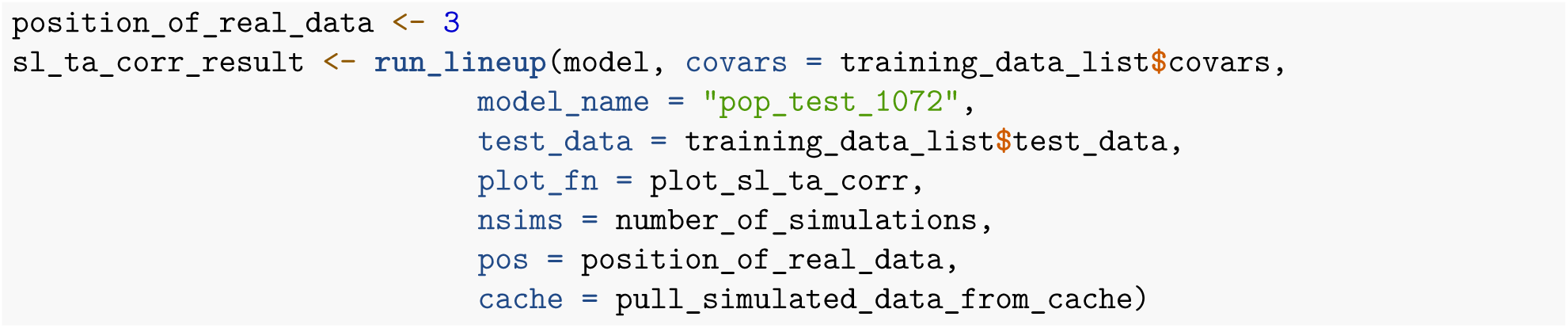

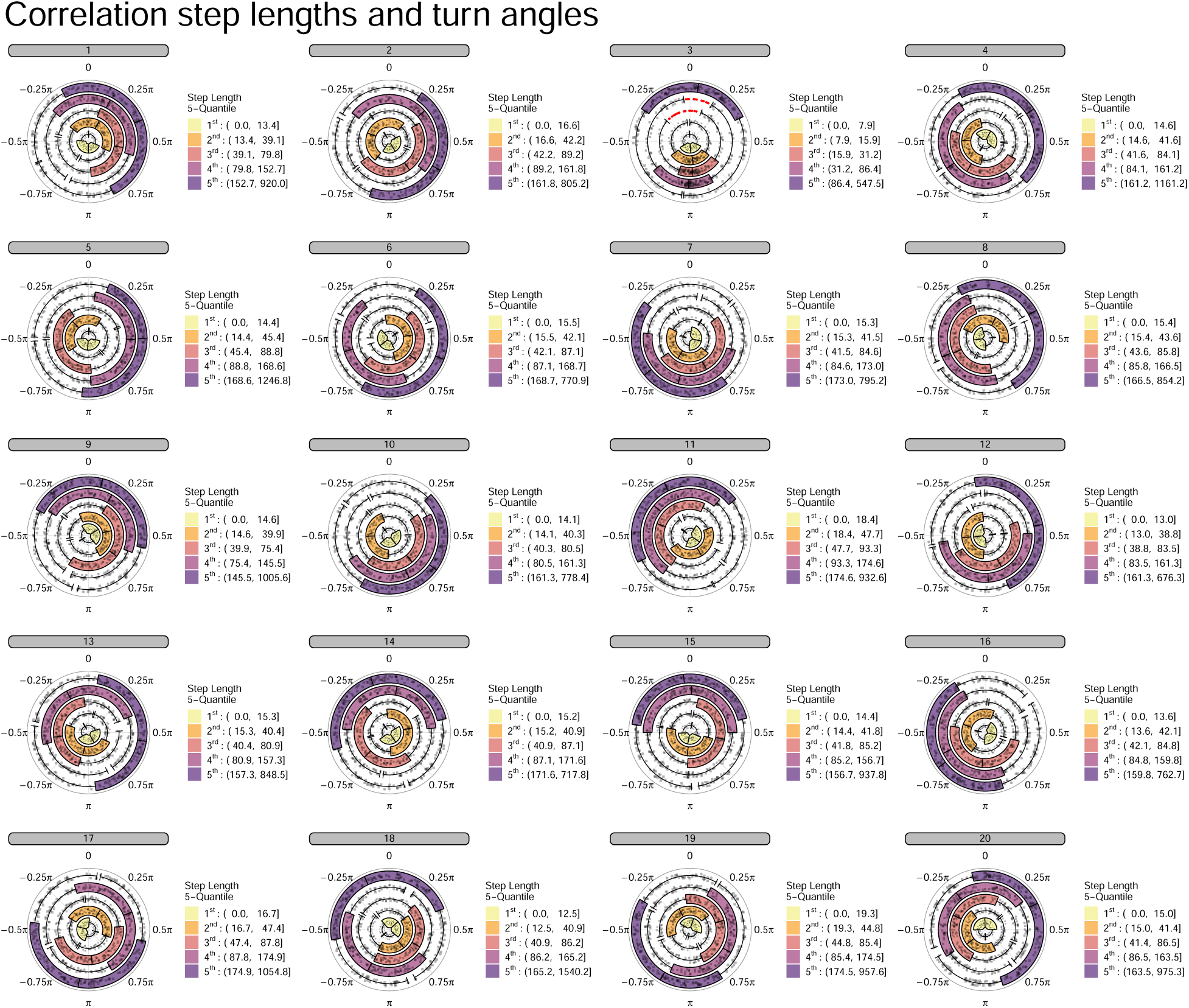

#### Document footer

##### Session Information

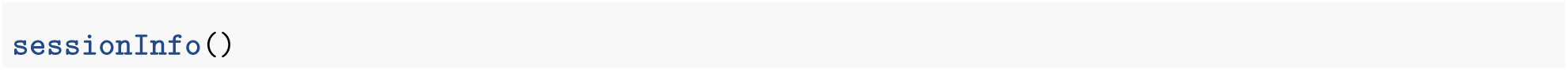

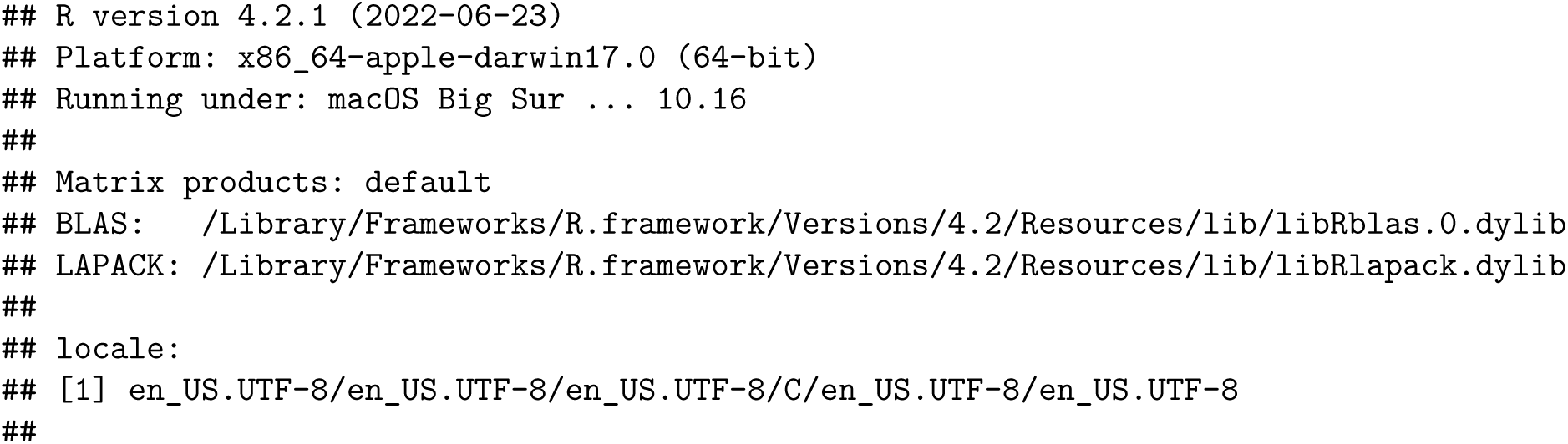
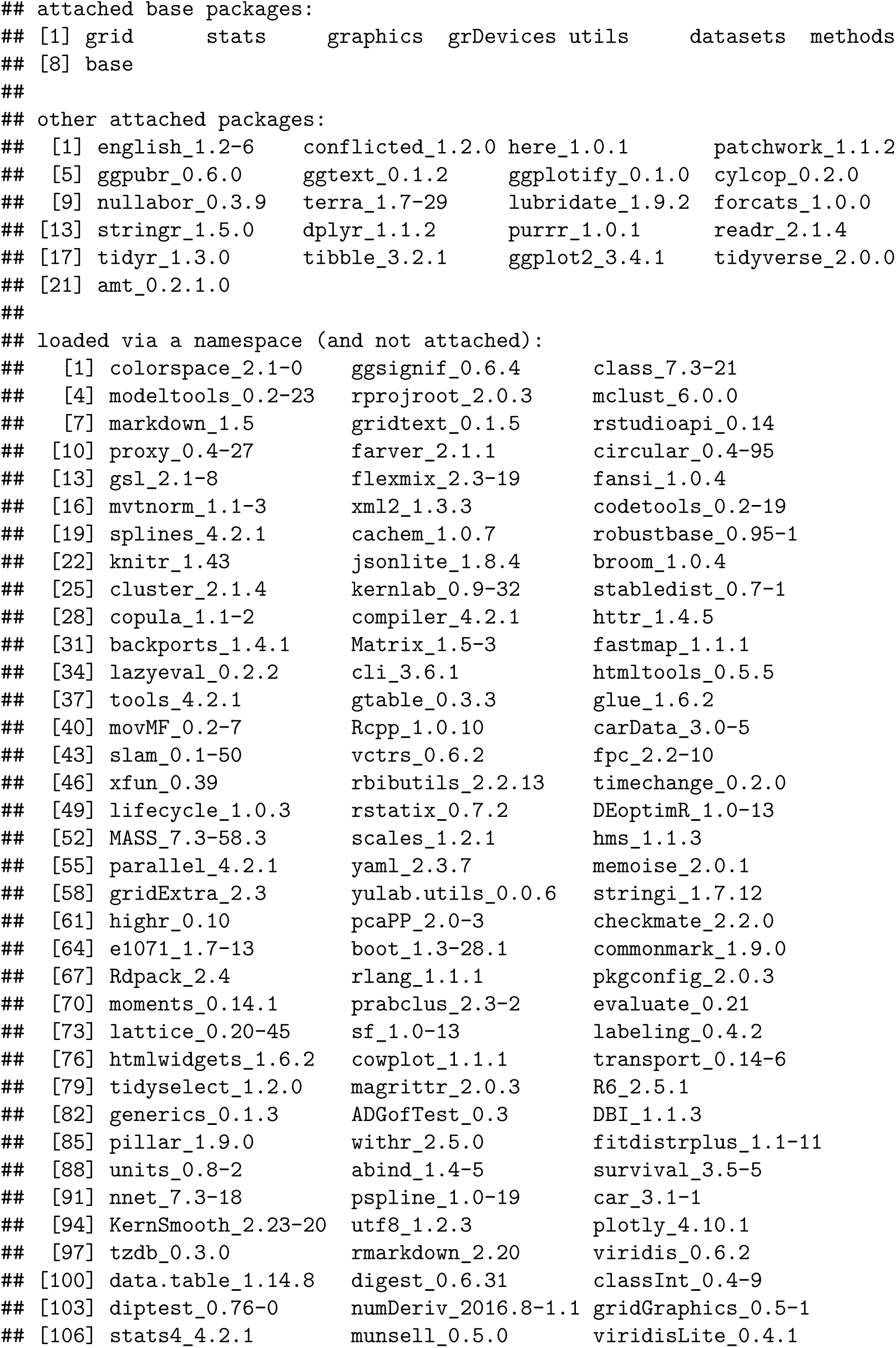

### Cross-validation of Animal Movement HMMs within the Lineup Protocol

09-22-2023

#### Document Preamble

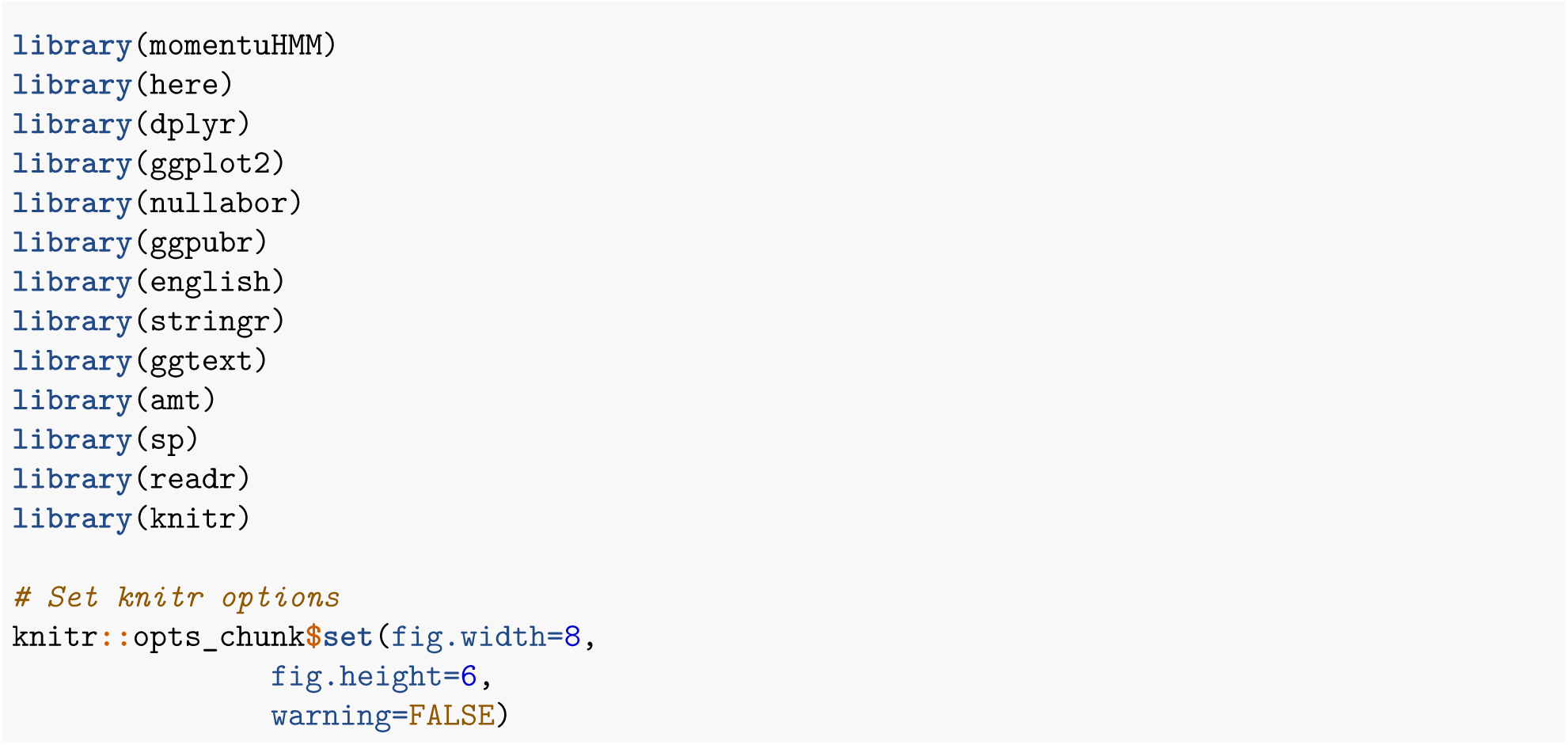

#### Intro

The purpose of this vignette is to demonstrate how to use the lineup protocol in tandem with animal movement hidden Markov models (HMMs) in order to do basic cross-validation of a given model. To illustrate how this process works, we’ve included the diagram below that includes specific references to functions defined later in the vignette. Though this diagram is somewhat tailored to our use-case, the steps are wildly applicable to other model types.

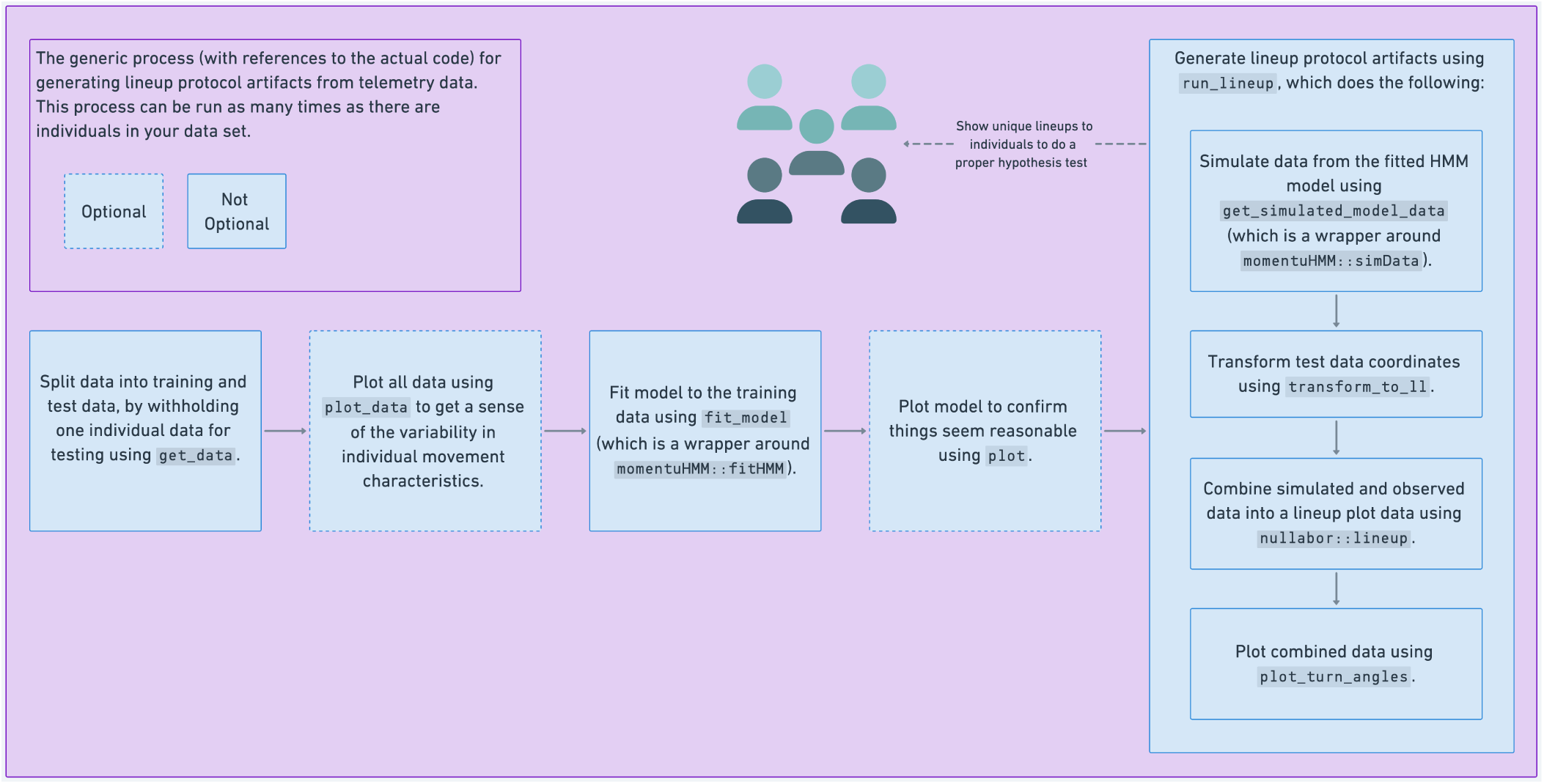

#### Training and Testing Data

In order to demonstrate a simplistic cross-validation approach using the lineups protocol, we have written the following function, get_data, which takes a given fisher ID whose data will be used to test (or cross-validate), the remaining data will be used to parameterize the model. We will fit the same model three times, each time retaining one of the fishers’ data for testing. This section also contains the function plot_data, which is a generic function for plotting movement characteristics (i.e., step-lengths, turn-angles, and trajectories).

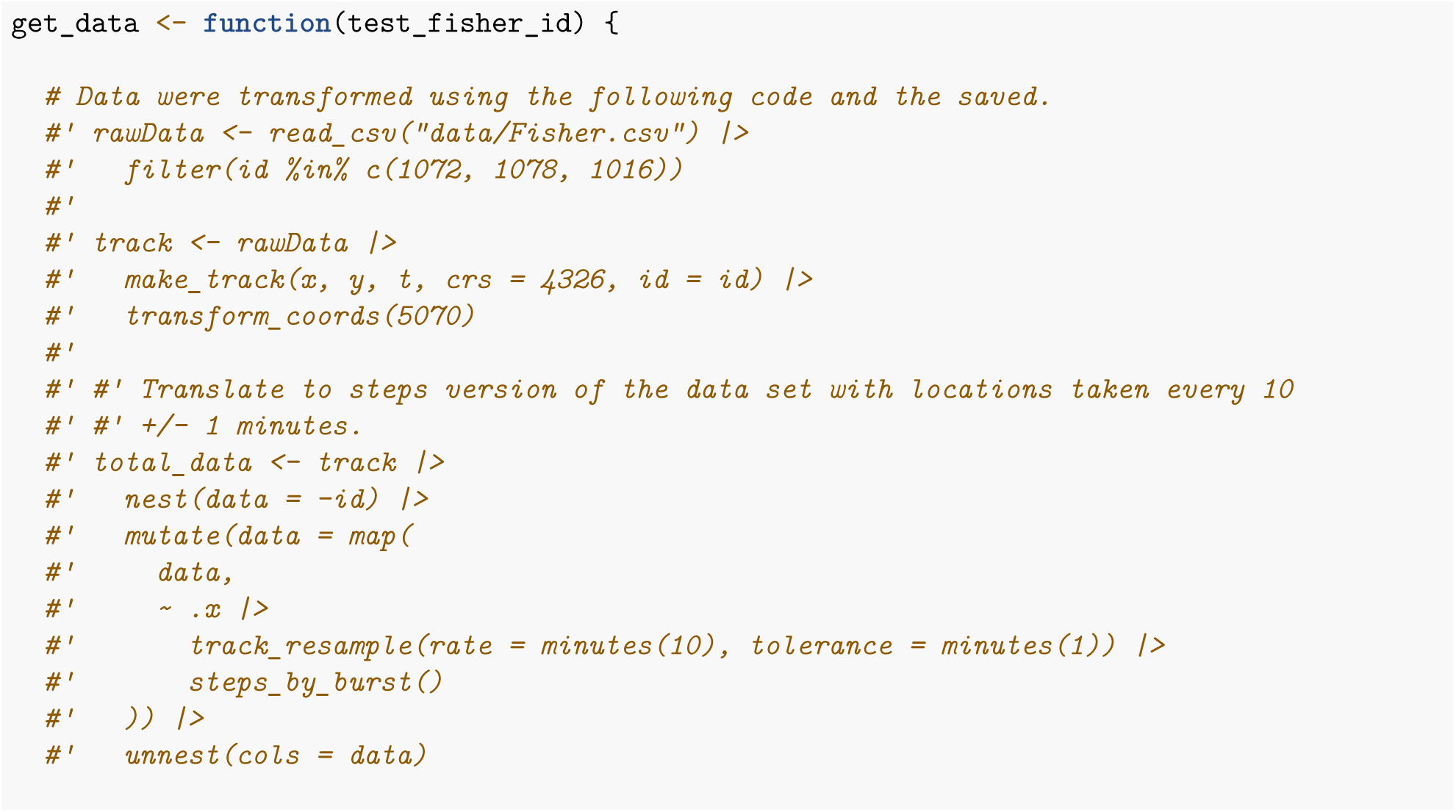
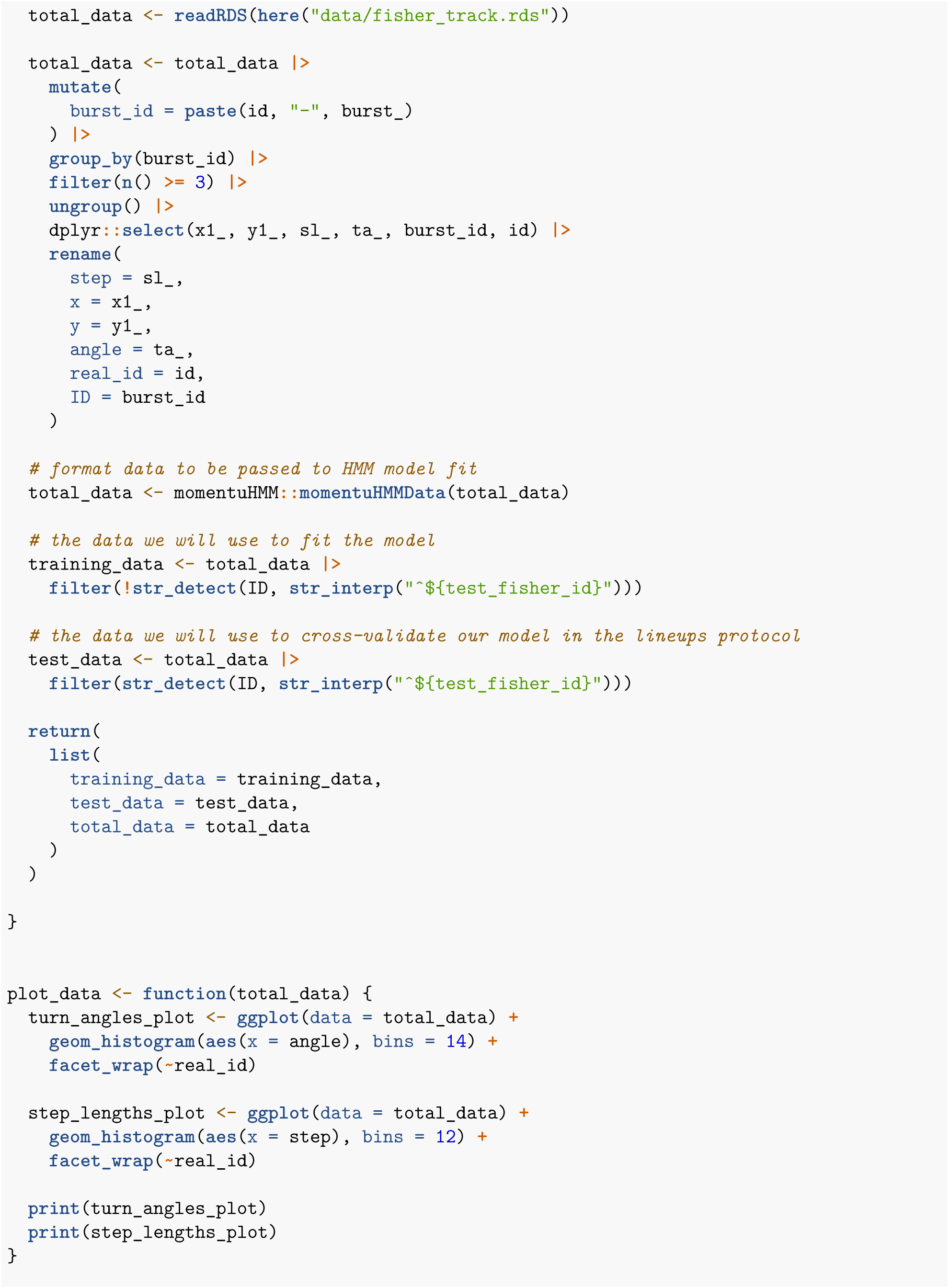
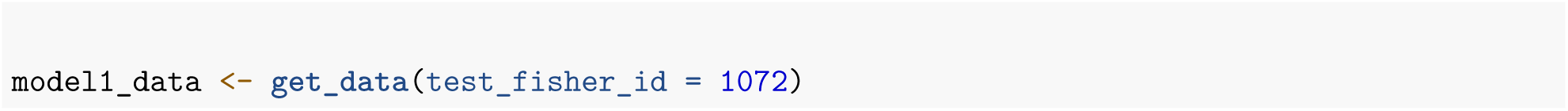

Let’s plot the total data to get a visual sense of how the three fishers’ movement differs from each other.

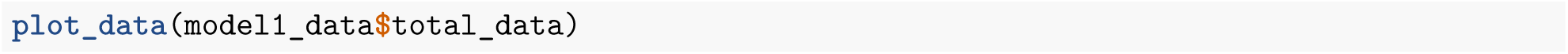

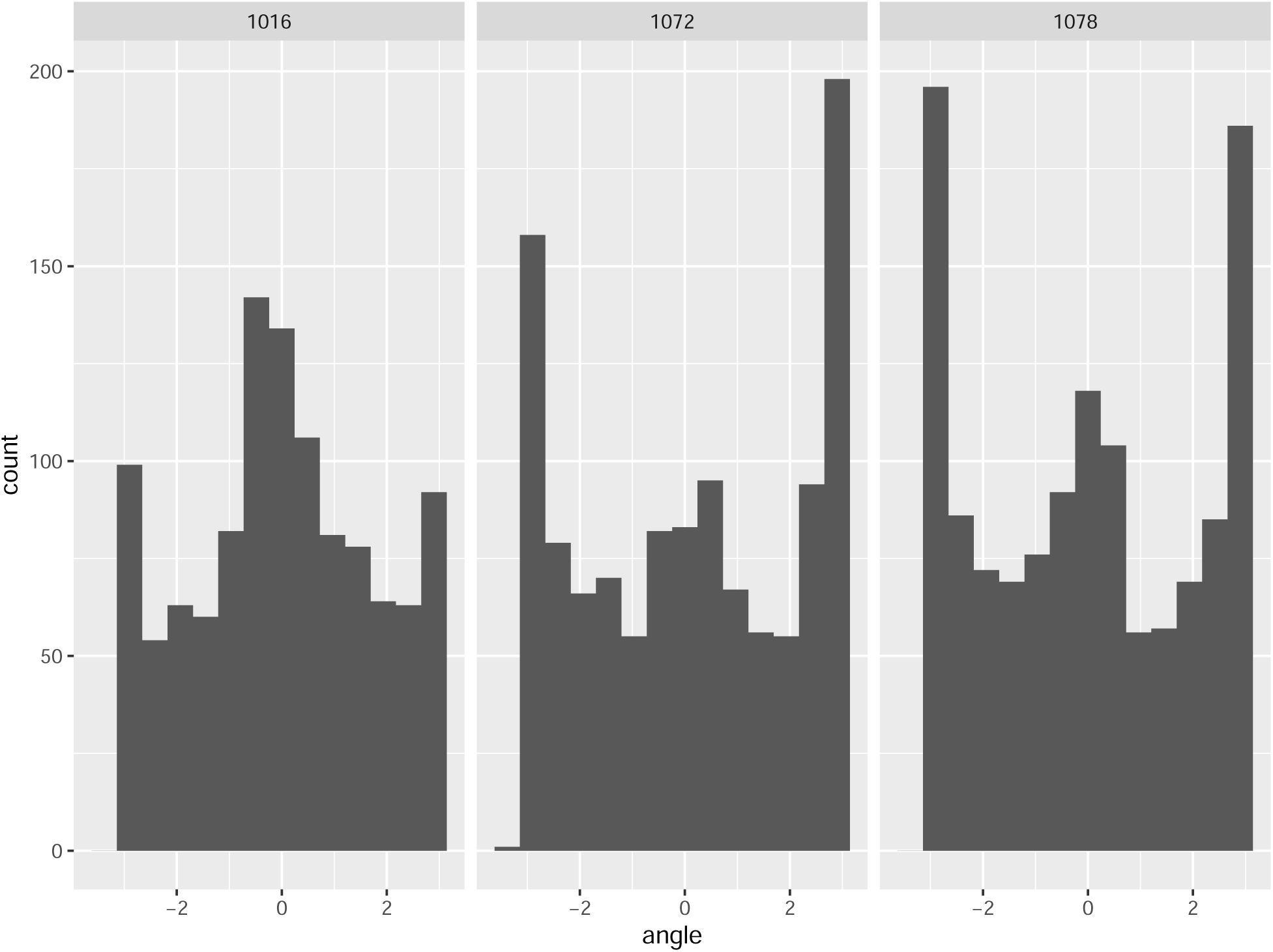

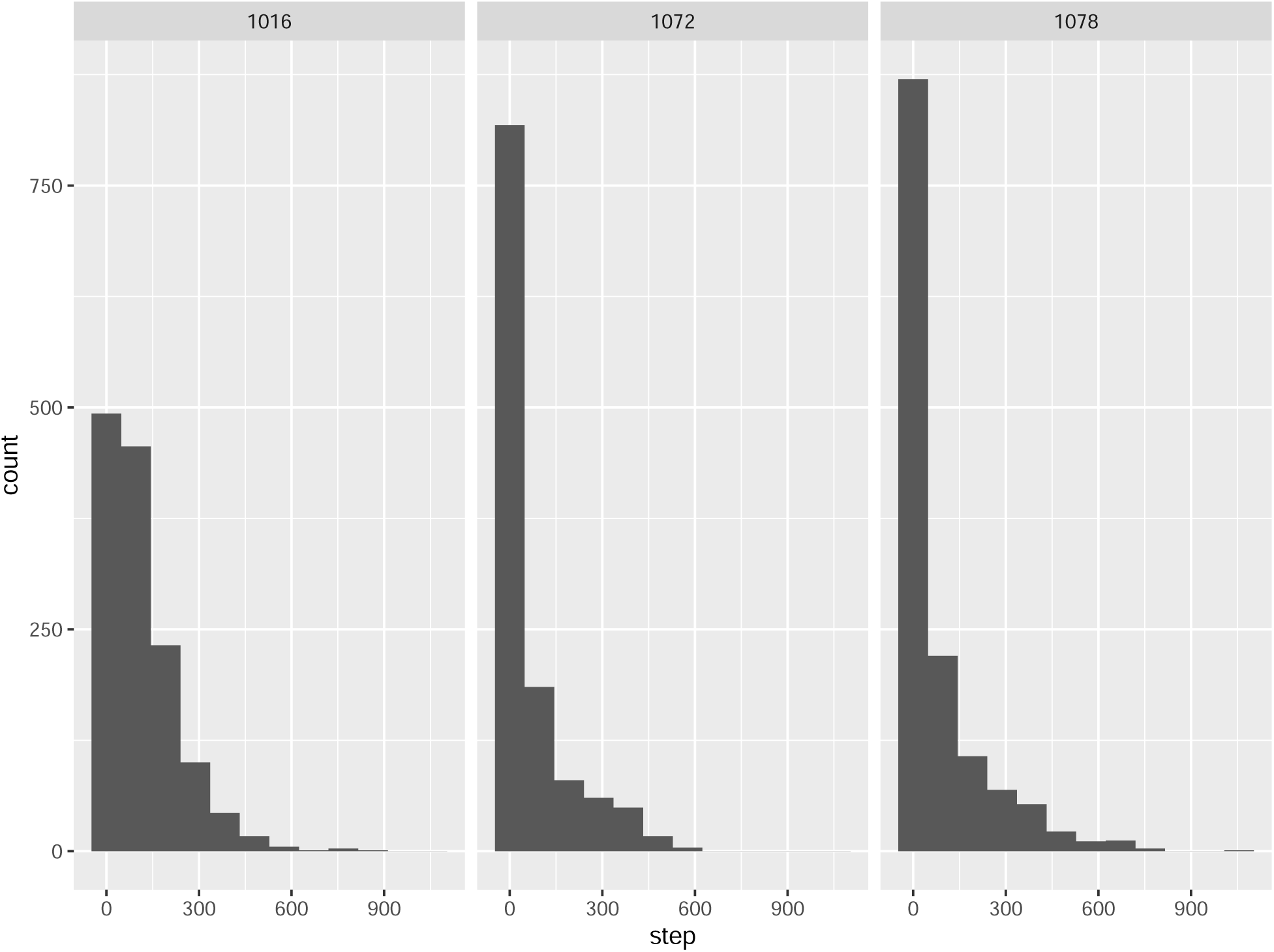

#### Fitting a Two-State HMM model

Now that we have our data, let’s actually fit the first model using our function fit_model and plot the model output.

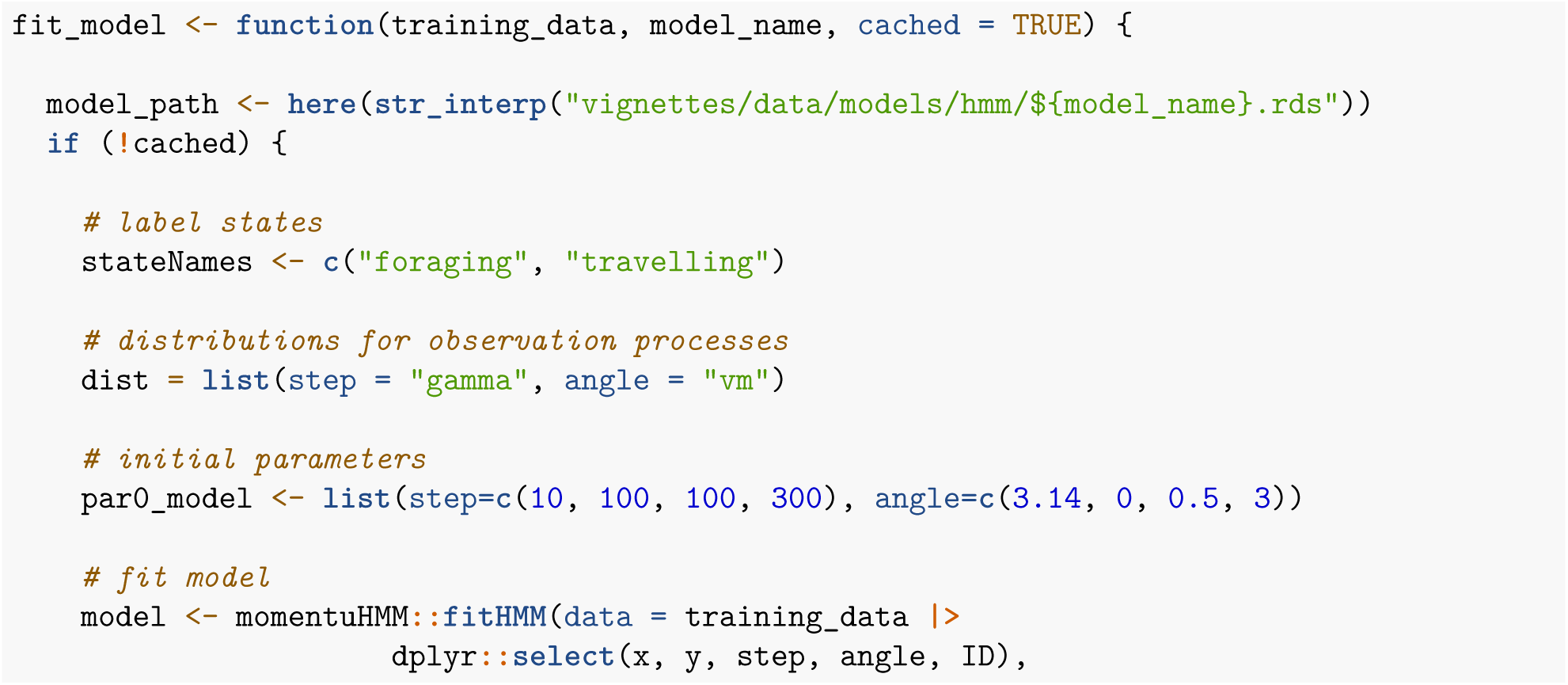
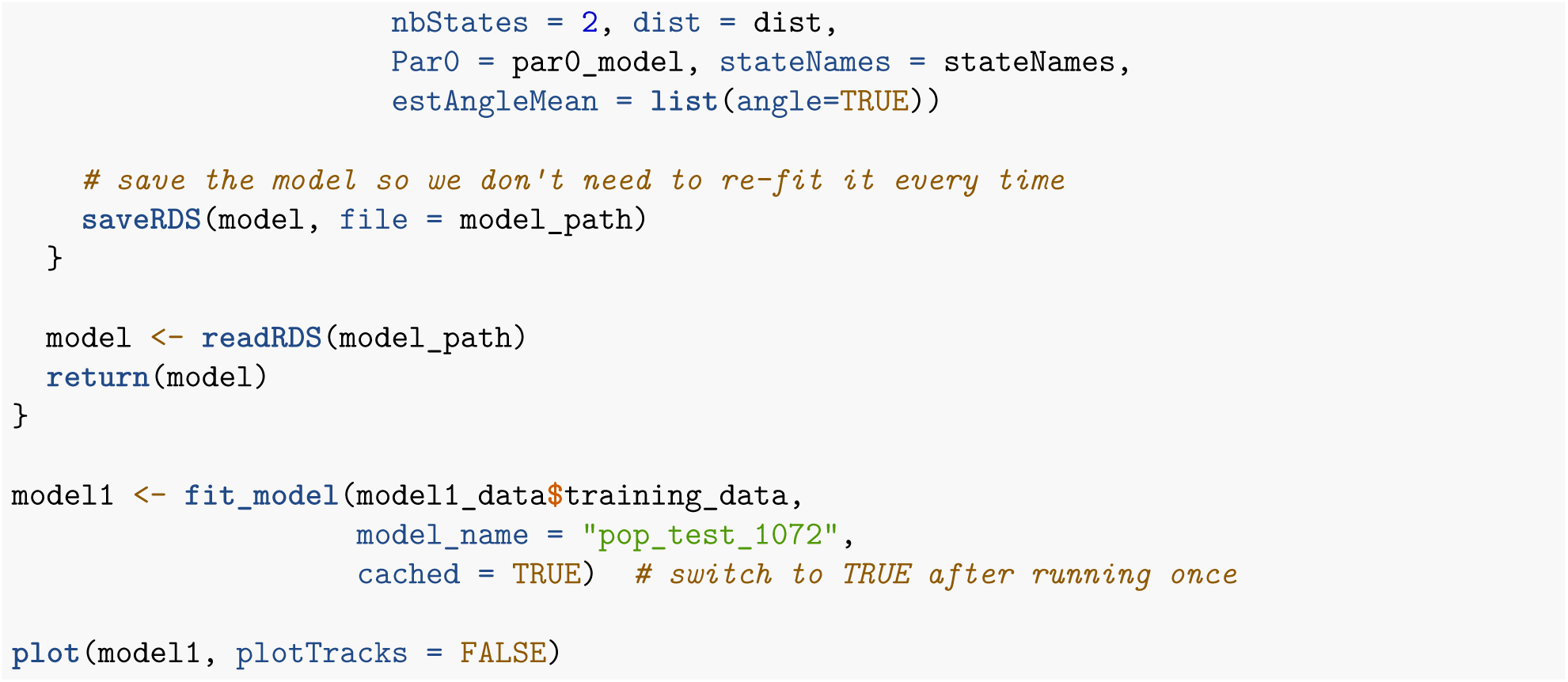

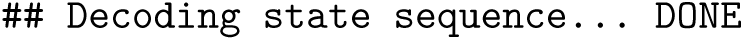

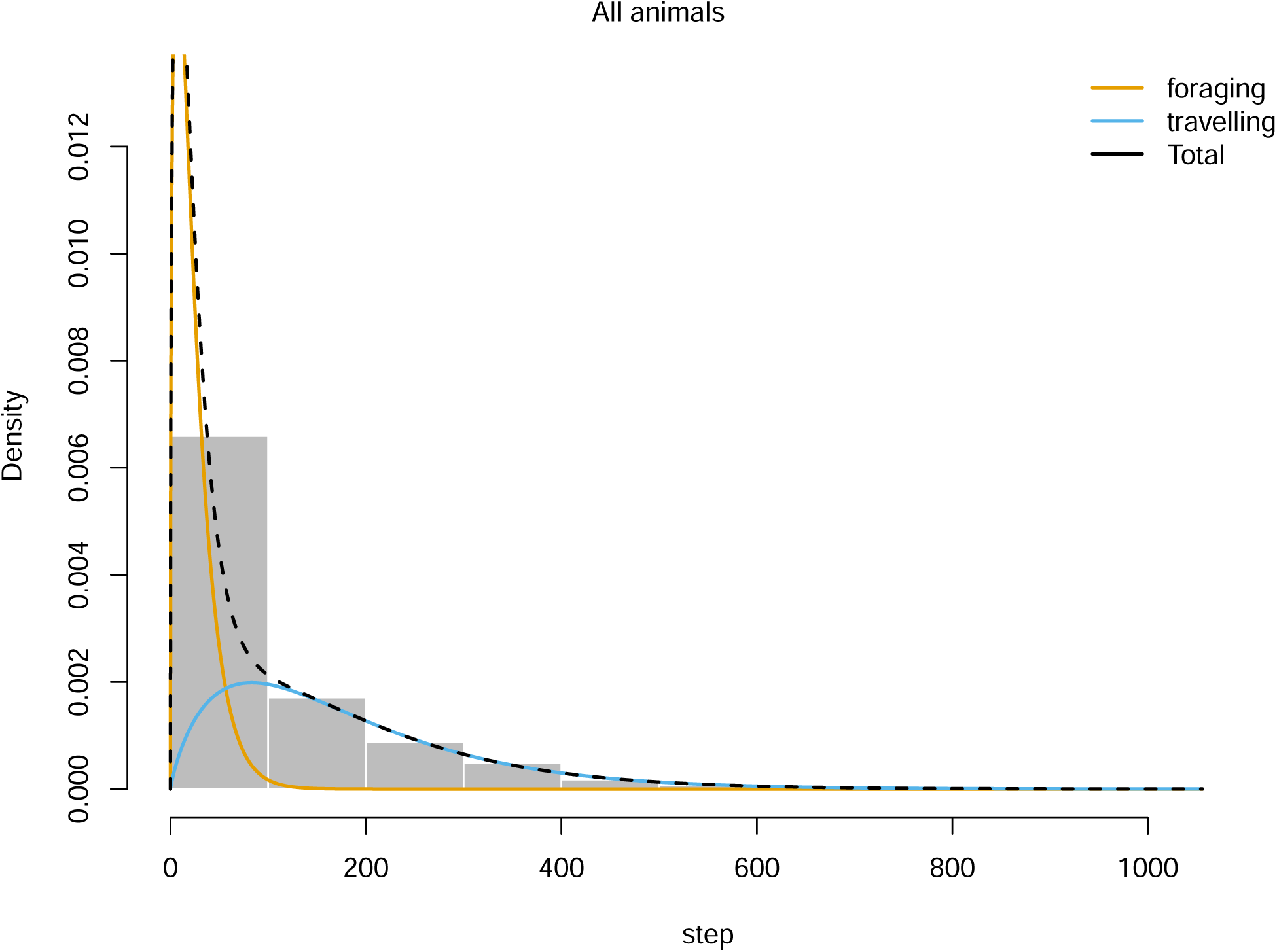

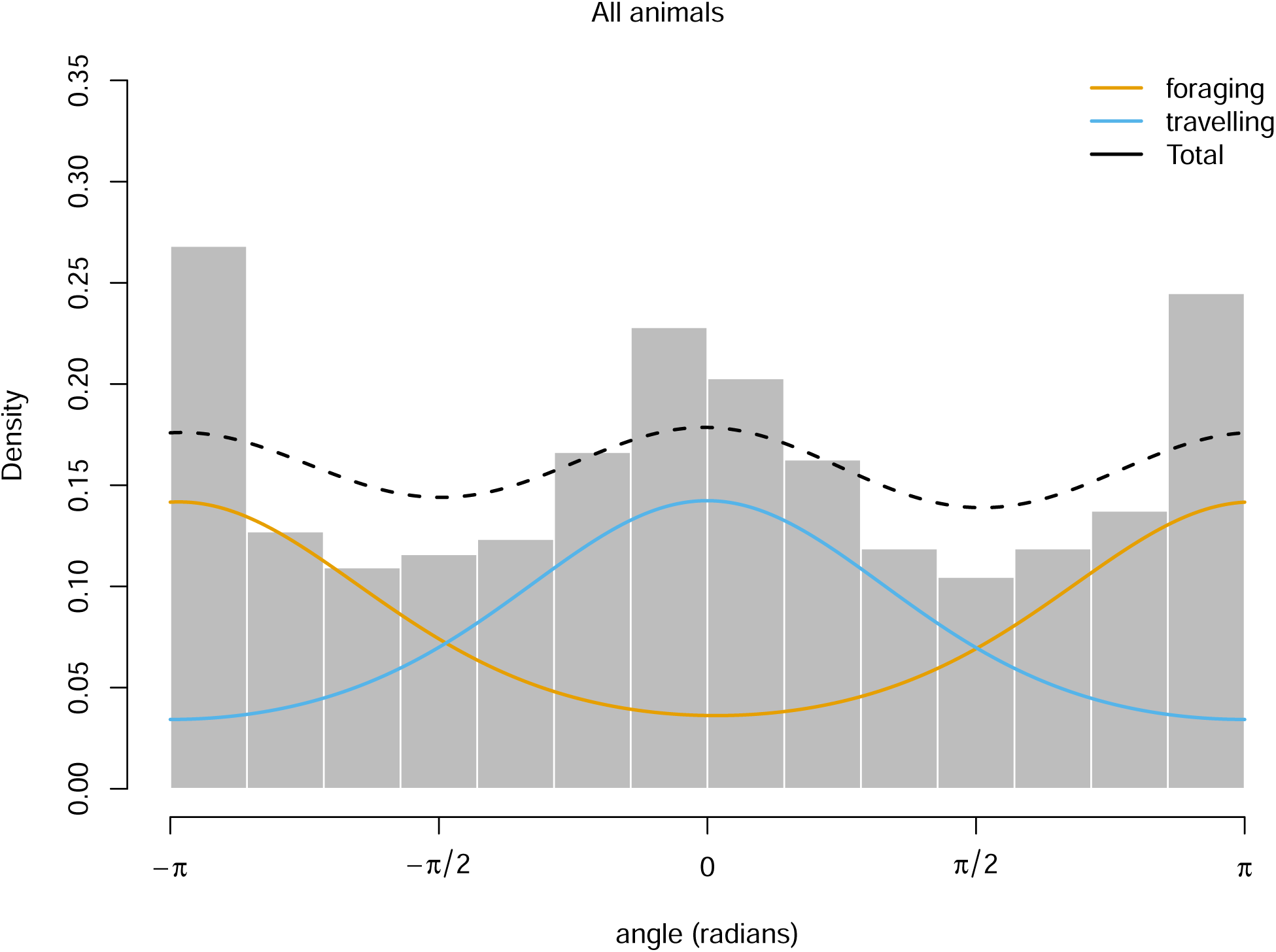

#### Generating Lineup Protocol Artifacts

For the sake of this vignette we will focus on creating turn-angle histograms. But remember! The lineup protocol can accommodate any visualization of the data.

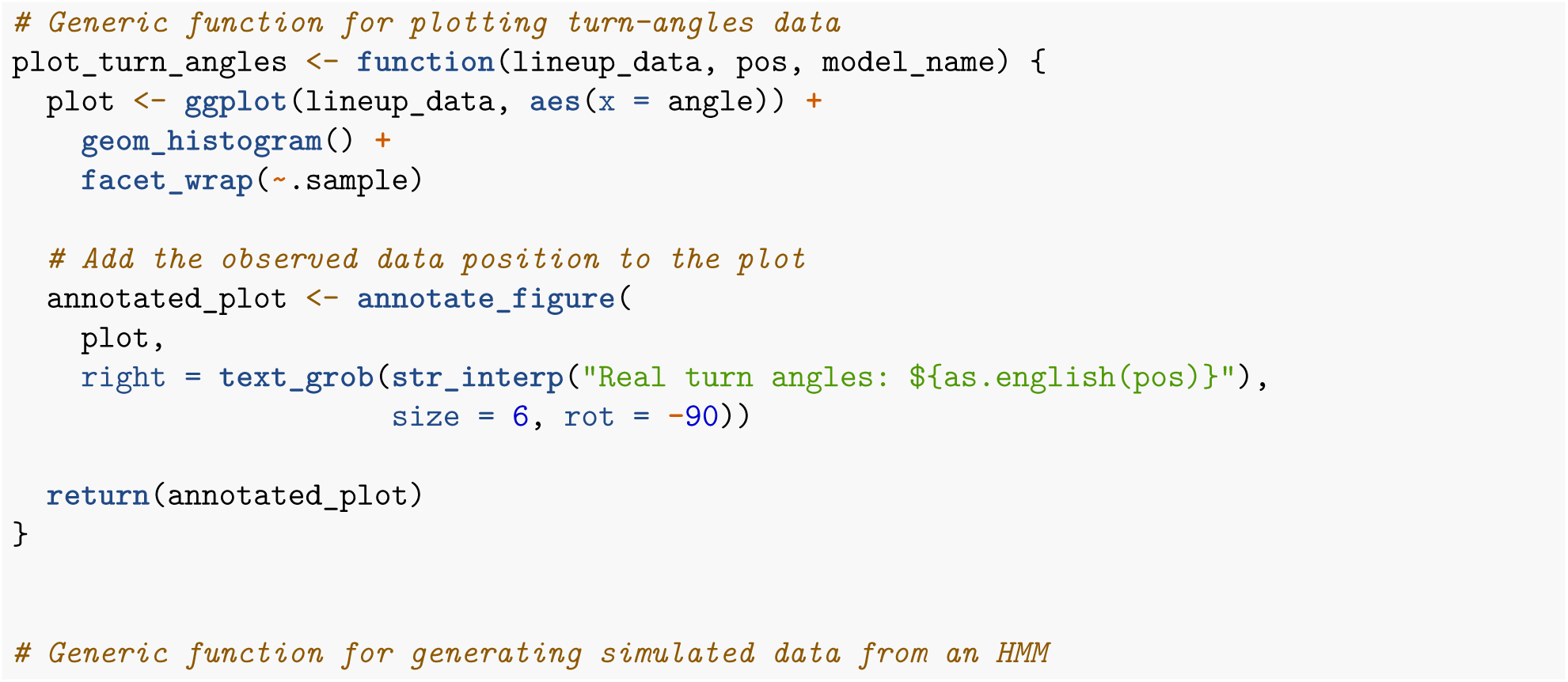
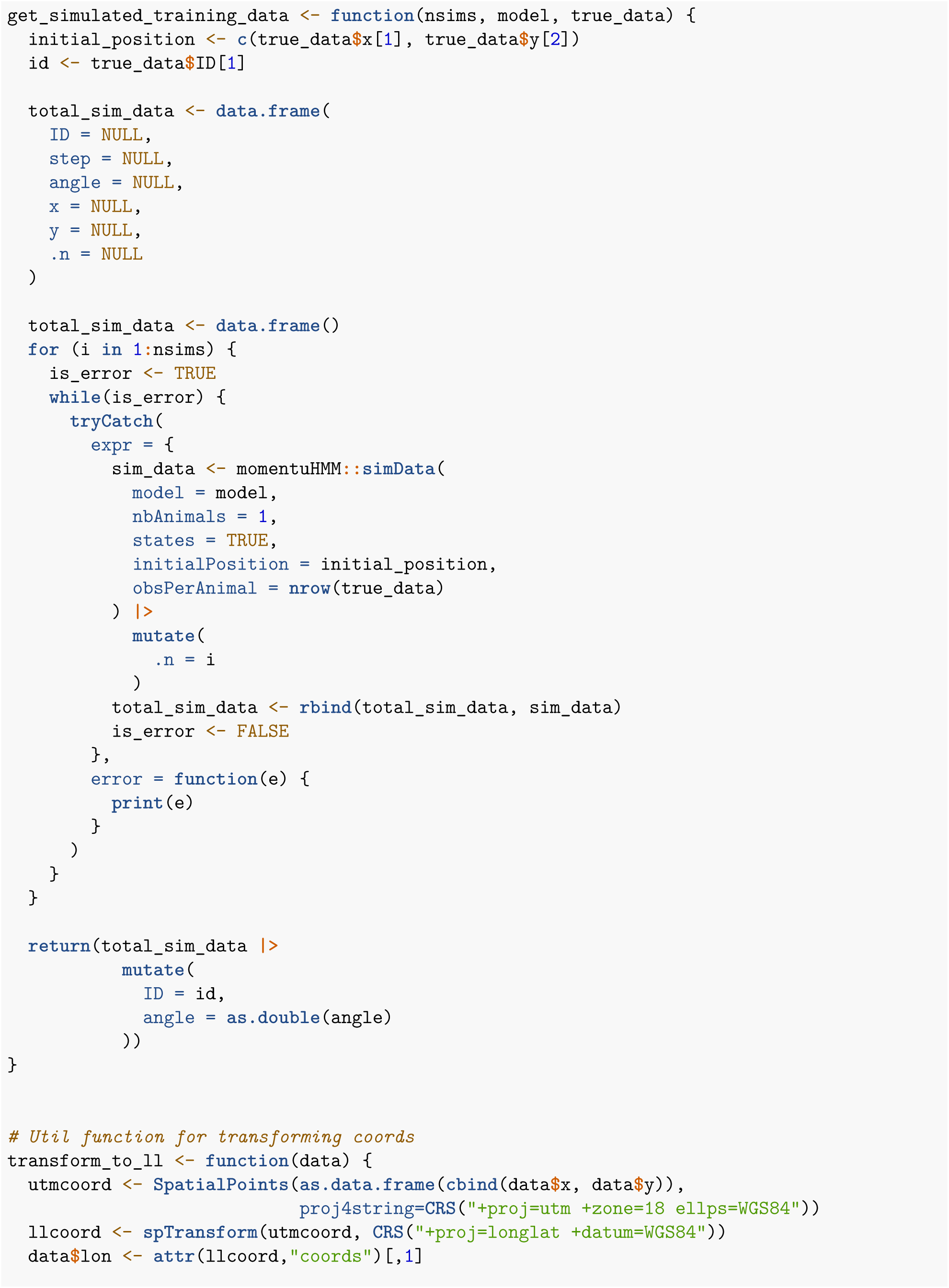
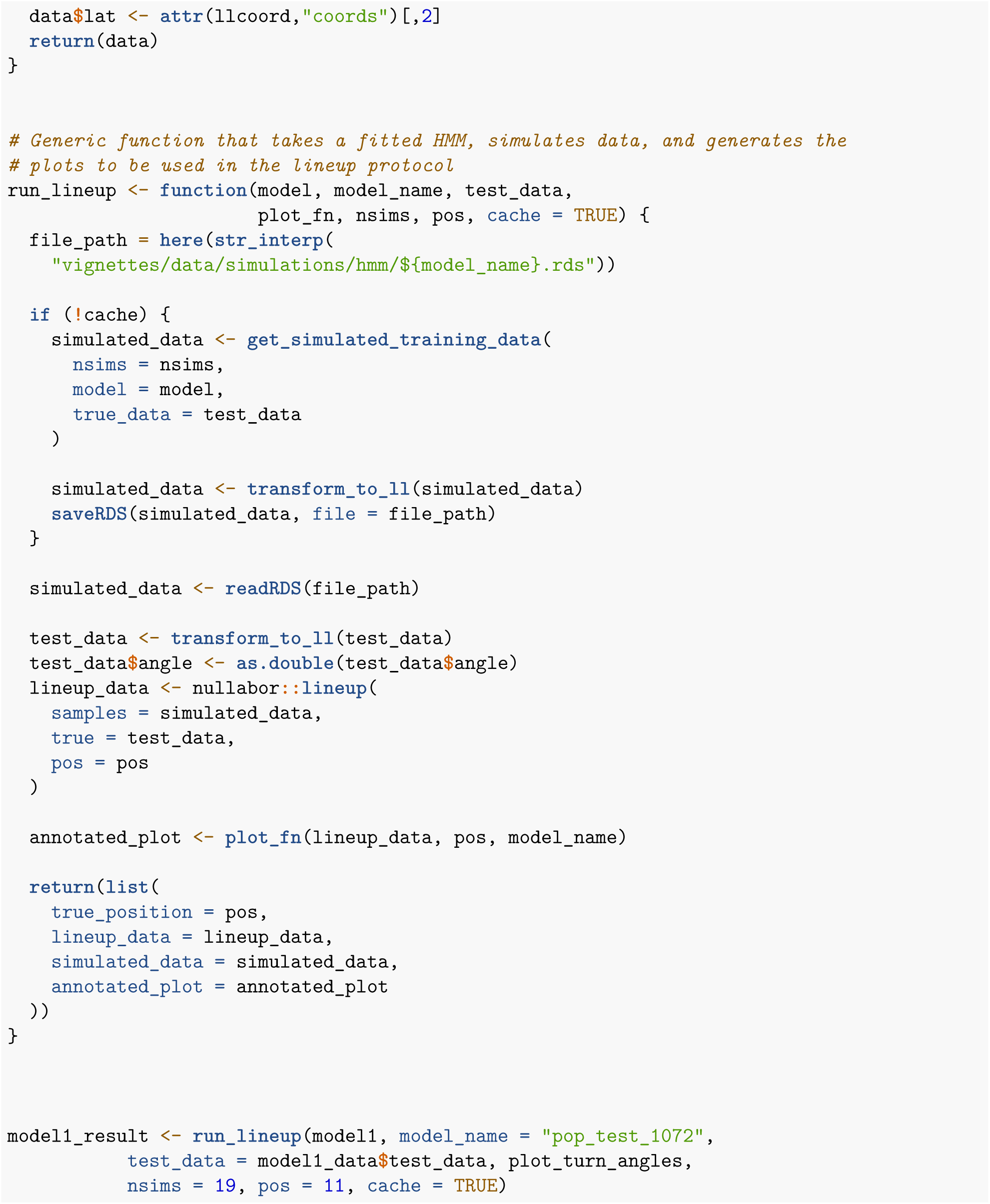
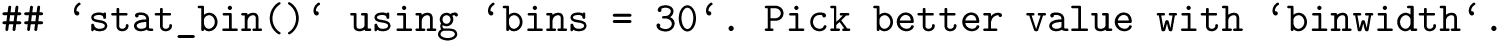

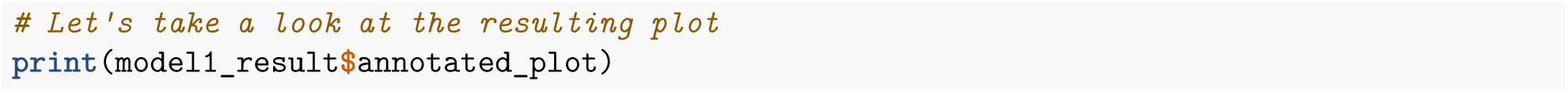

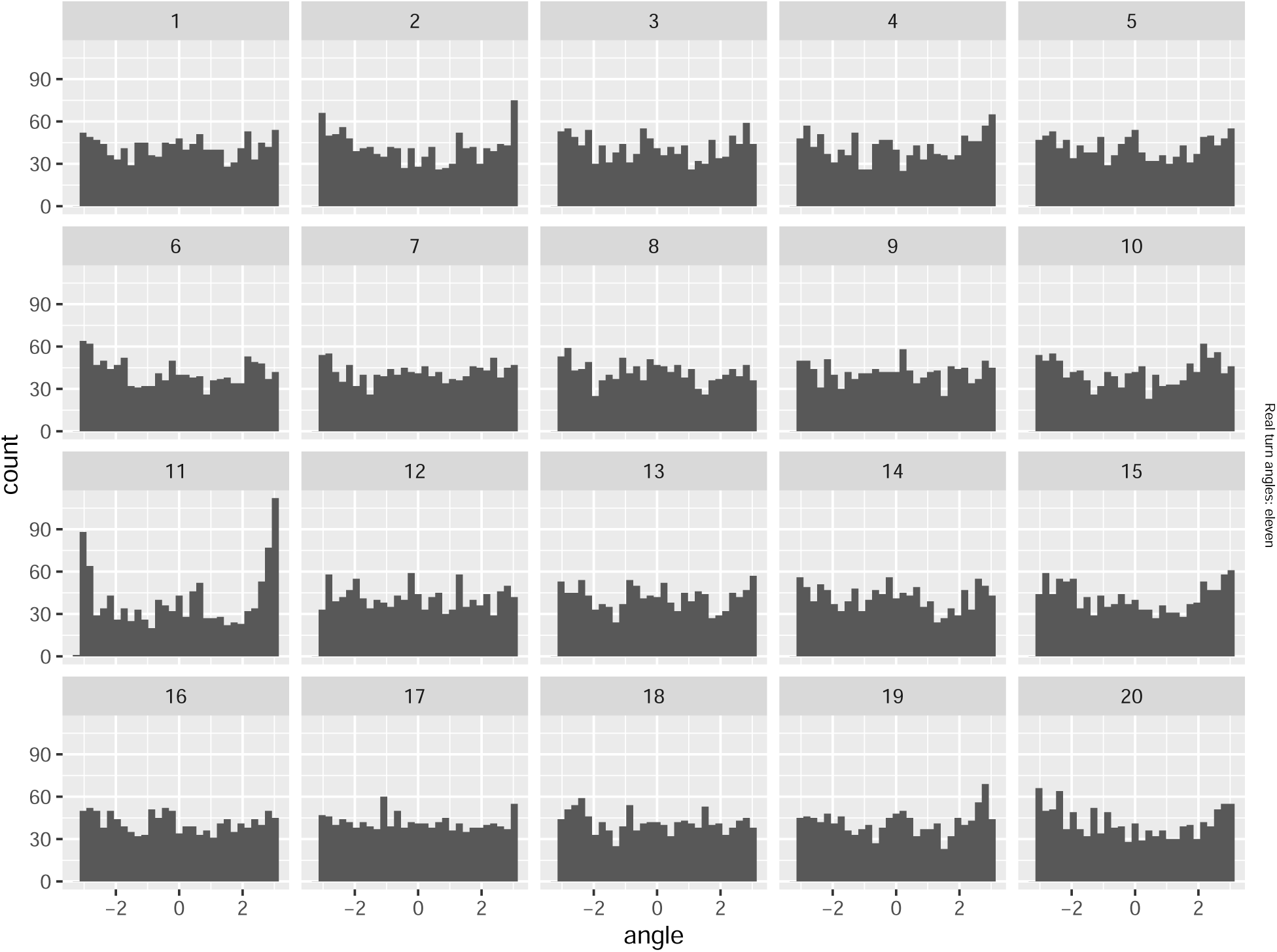

Now that we’ve generated a lineup protocol artifact for a set of data, let’s do the same for the remaining two sets. This time we will reserve data from fisher 1078 for testing and execute all the same code we did for our first iteration.

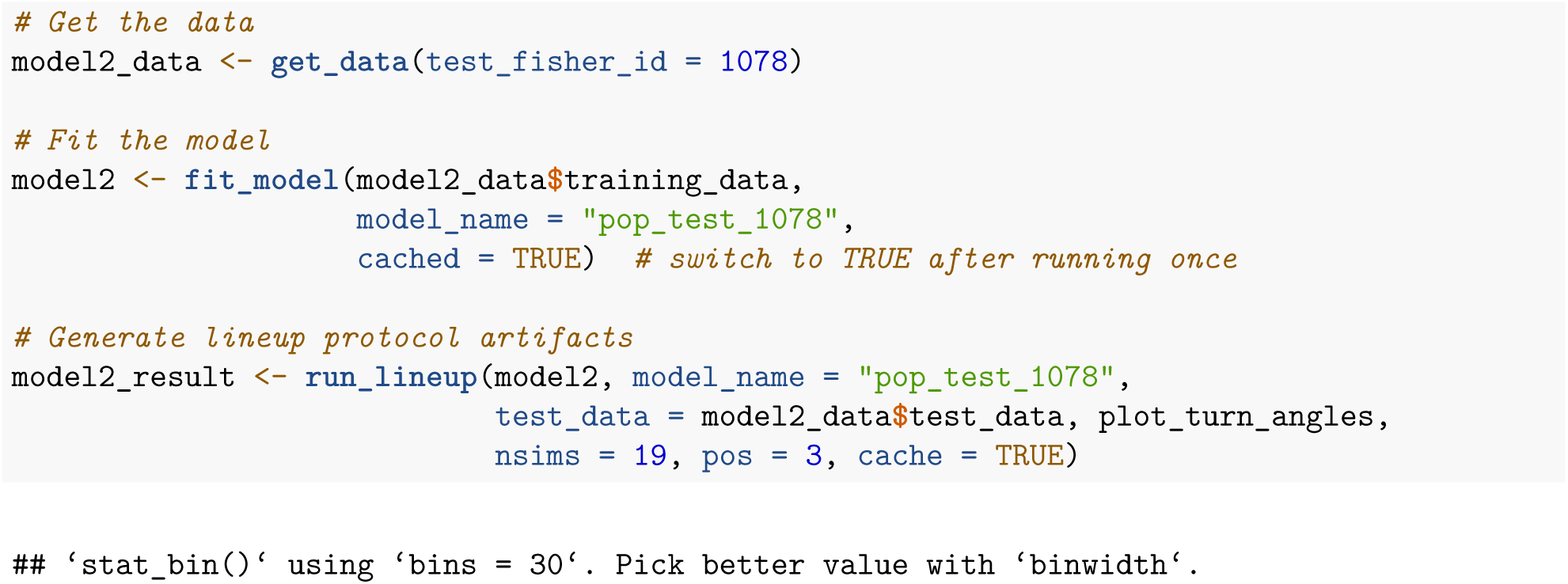
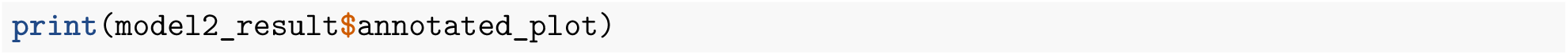

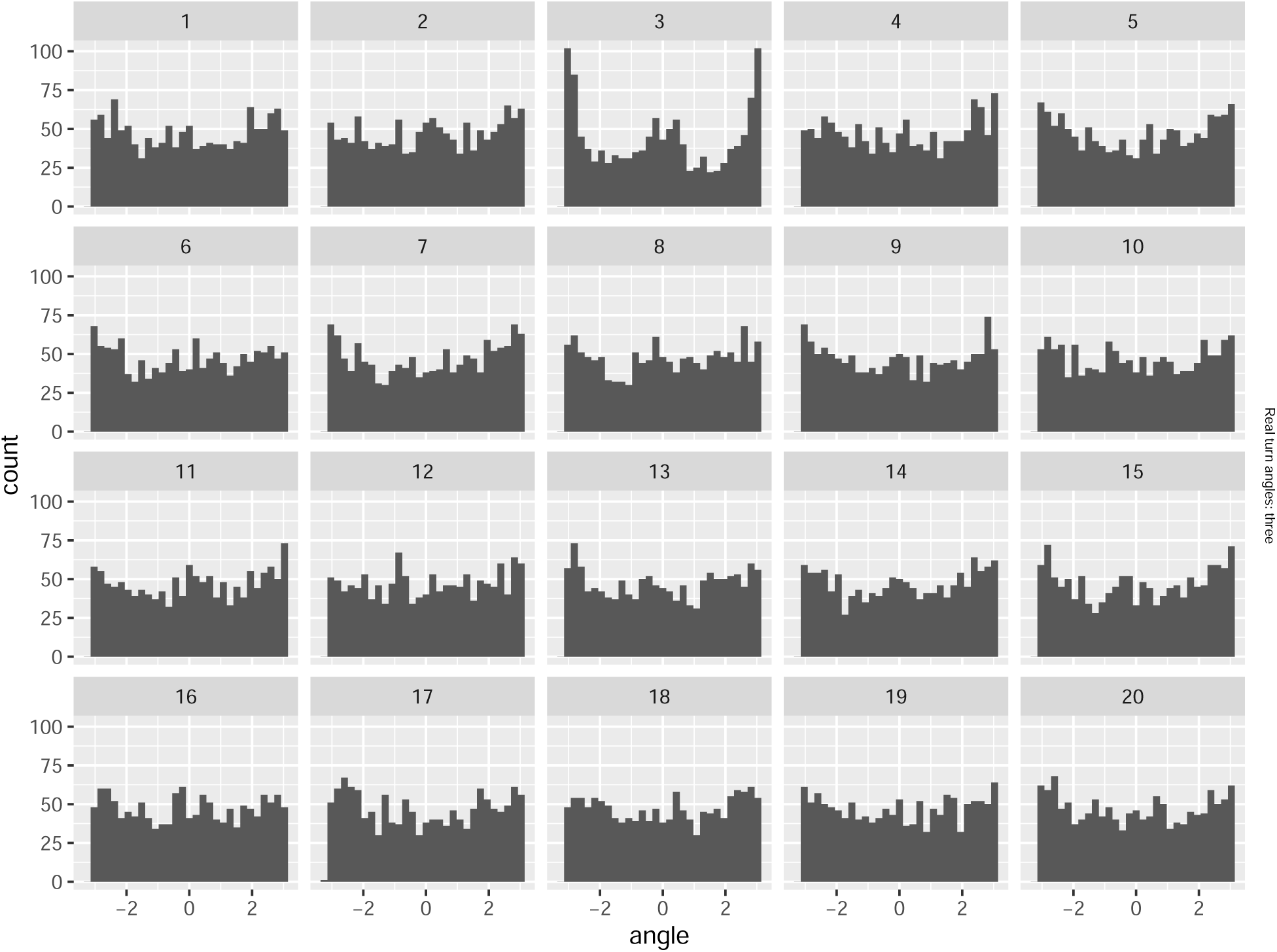

Alright, let’s fit our final model, this time reserving fisher 1016 for testing

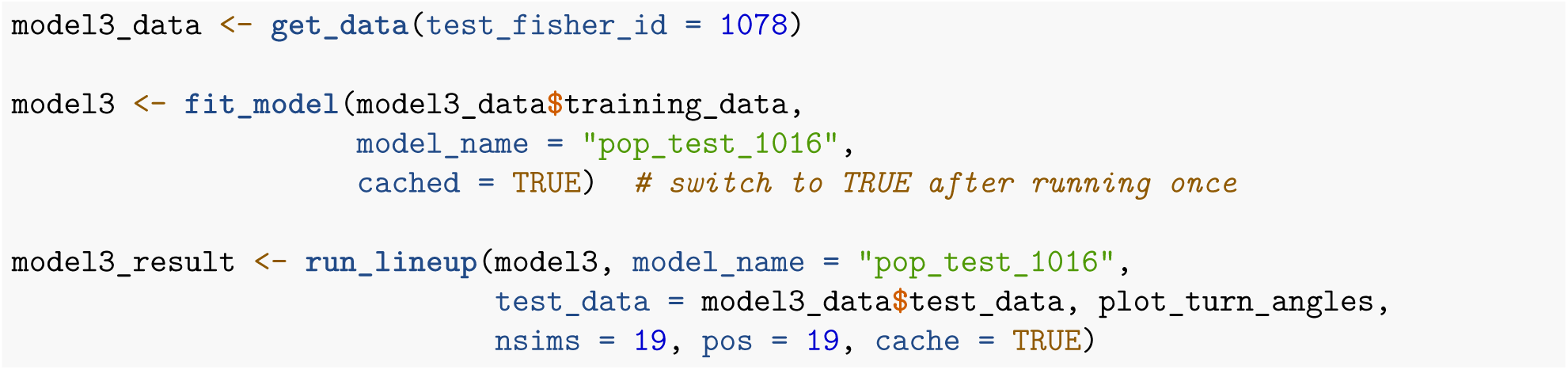

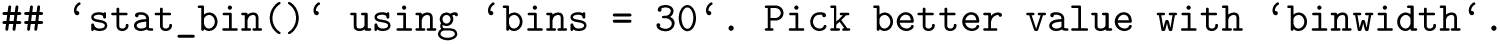

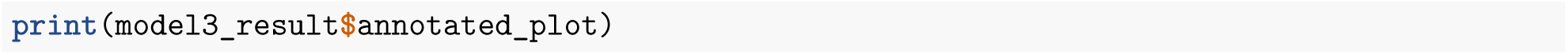

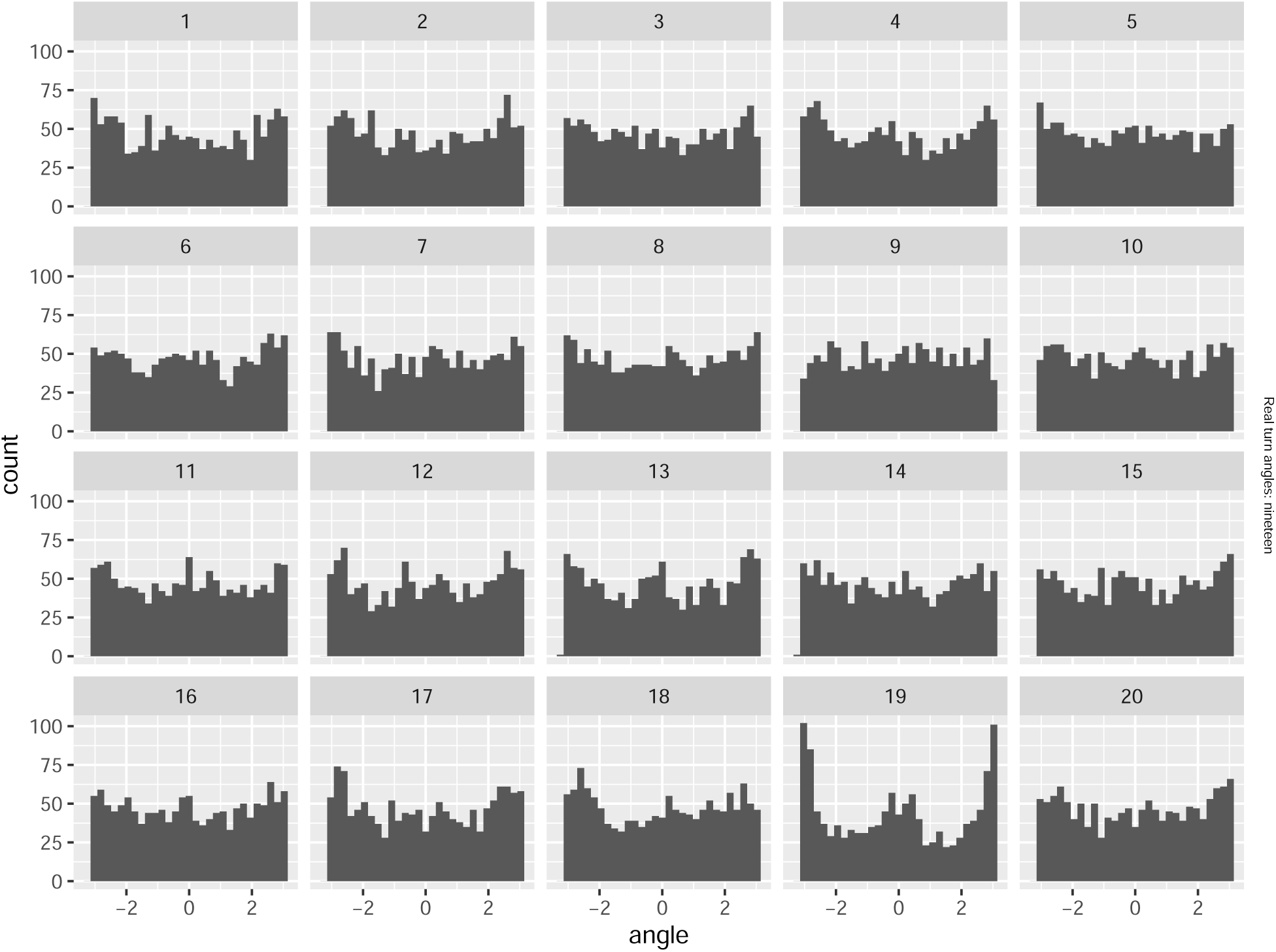

#### Document footer

##### Session Information

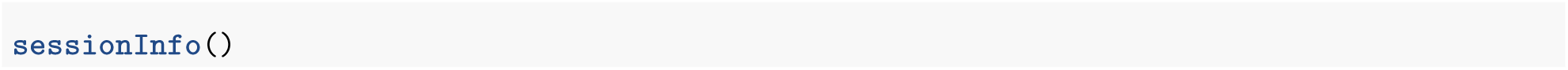

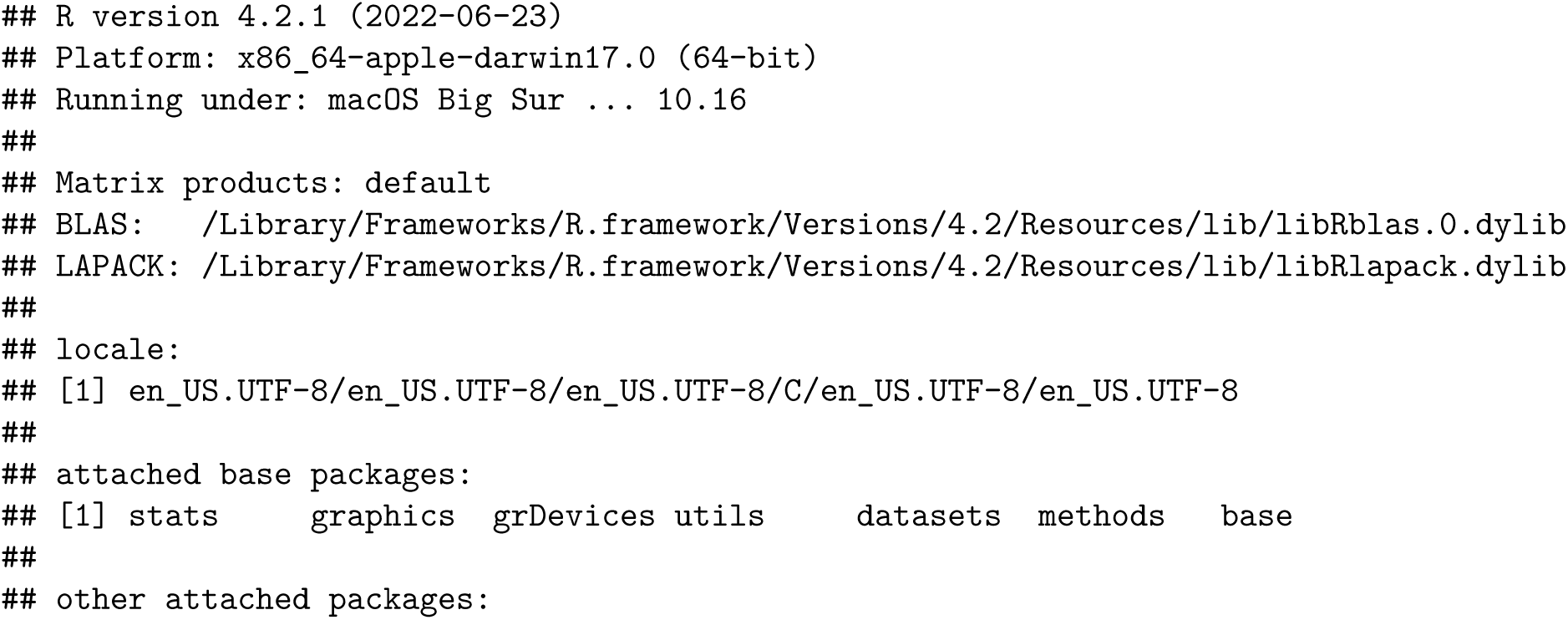
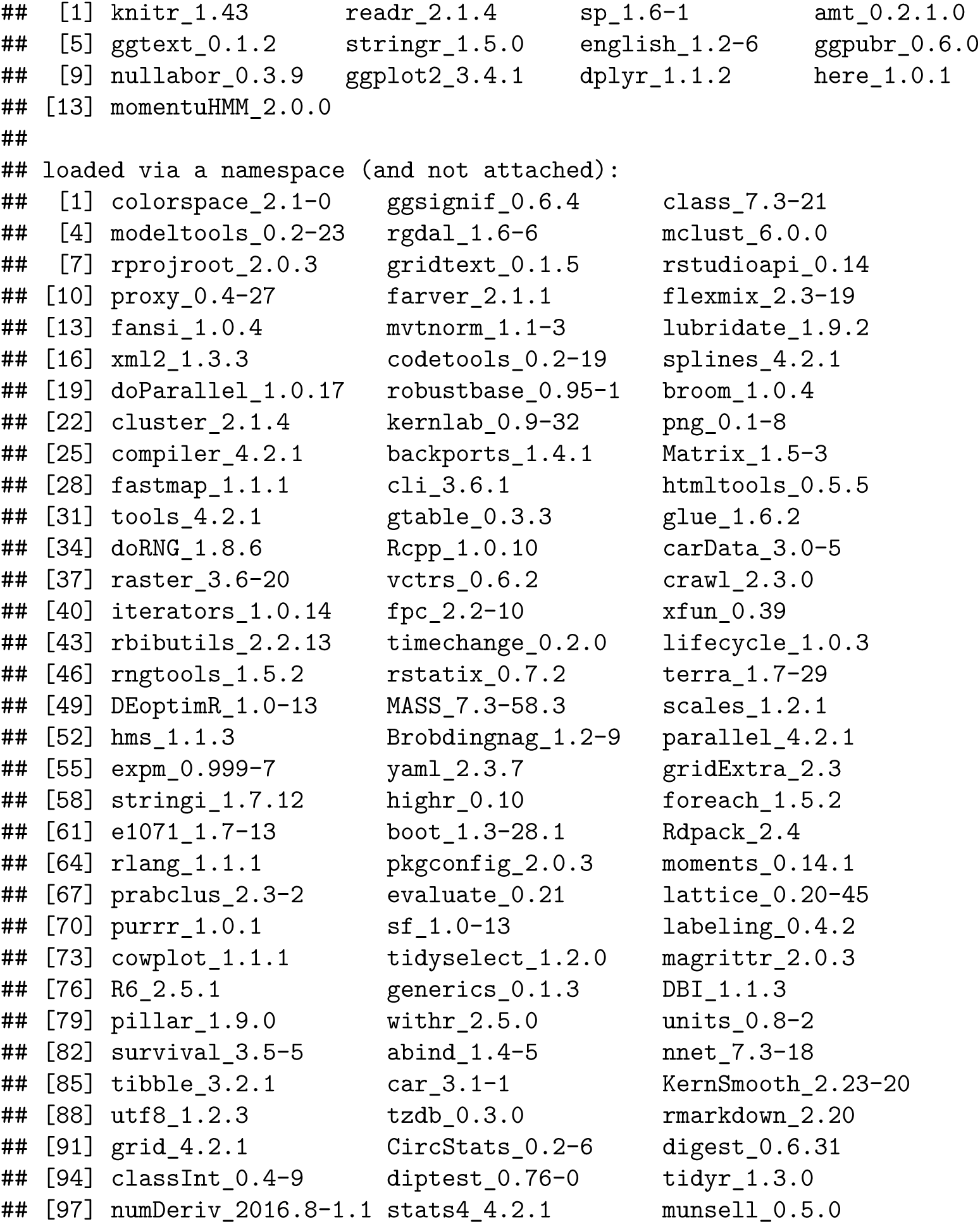

